# Accurate, scalable cohort variant calls using DeepVariant and GLnexus

**DOI:** 10.1101/2020.02.10.942086

**Authors:** Taedong Yun, Helen Li, Pi-Chuan Chang, Michael F. Lin, Andrew Carroll, Cory Y. McLean

## Abstract

Population-scale sequenced cohorts are foundational resources for genetic analyses, but processing raw reads into analysis-ready variants remains challenging. Here we introduce an open-source cohort variant-calling method using the highly-accurate caller DeepVariant and scalable merging tool GLnexus. We optimized callset quality based on benchmark samples and Mendelian consistency across many sample sizes and sequencing specifications, resulting in substantial quality improvements and cost savings over existing best practices. We further evaluated our pipeline in the 1000 Genomes Project (1KGP) samples, showing superior quality metrics and imputation performance. We publicly release the 1KGP callset to foster development of broad studies of genetic variation.

## Background

Sequencing a single individual can identify variants informative for diseases [1], traits [2], and ancestry [3]. Jointly using sequence data of multiple individuals can discover rare genetic diseases [4]. Population-scale sequencing generates annotation resources for clinical sequencing, such as dbSNP [5], ExAC [6], DiscovEHR [7], TOPMed [8], and gnomAD [9], and enables well-powered association studies [10] in large datasets of sequenced and phenotyped individuals [11].

Single-sample variant calling methods [12–16] use sequence reads mapped to a reference genome to identify and genotype positions which differ from the reference. Many variant callers support the generation of Genome Variant Call Format (gVCF) outputs, which supplement the variant calls with block records of non-variant regions annotated with confidence estimates that the regions match the reference genome. Joint genotyping tools such as GATK GenotypeGVCFs [17] and GLnexus [18] transform a cohort of gVCFs into a project-level VCF that contains a complete matrix of every variant in a cohort with a call for each sample. Compared to a full joint-calling strategy, joint genotyping both substantially reduces the size of required input data and avoids the need to fully reprocess all samples when adding samples to an existing cohort.

Joint genotyping of large cohorts introduces unique challenges. Harmonizing the representation of overlapping alleles is algorithmically intricate, and the number of overlapping alleles increases with cohort size. In addition, even with high single-sample variant calling accuracy, many samples will aggregate a large number of total errors. At the same time, large cohorts present unique opportunities to increase accuracy. Cross-referencing genotype likelihoods across a cohort can help refine calls and filter errors, for example by identifying recurrent artifacts that violate Hardy-Weinberg equilibrium (HWE) [19].

Here we introduce a framework to generate highly accurate and scalable cohort callsets with DeepVariant, using its superior calibration of variant confidences and high single-sample accuracy [15]. We adapt the scalable joint genotyper GLnexus [18] to DeepVariant gVCFs and tune filtering and genotyping parameters to optimize performance for whole-genome sequences (WGS) and whole-exome sequences (WES) across a range of sequence coverages and cohort sizes. We compare the resulting callsets to analogous callsets generated by the broadly-used GATK Best Practices [20] which serve as current state-of-the-art benchmarks. Finally, we apply the optimized method to the recent deep sequencing of the 1000 Genomes Project (1KGP) phase 3 samples [21]. We evaluate the resulting callset across multiple quality metrics and performance as an imputation reference panel against a callset independently generated using a GATK Best Practices pipeline.

## Results

### Cohort variant call evaluation strategies

Four different measures of variant calling accuracy were used to optimize and evaluate cohort variant calls. First, we computed concordance of the GIAB HG001-HG007 benchmark samples to directly measure variant accuracy within the well-characterized 83% of the genome in the GRCh38 GIAB v3.3.2 benchmark regions. Second, we computed Mendelian violation rates in trios to indirectly measure variant accuracy genome-wide. Third, we computed the Ti:Tv ratio of single nucleotide polymorphisms (SNPs) for all samples to measure deviations from the expected genome-wide ratio of ~2.0-2.1 [22]. Fourth, we computed deviations from HWE at the cohort level, which can be indicative of recurrent artifacts in a variant calling algorithm [23].

### Cohorts used in development and evaluation

Four distinct data sources were used to optimize and evaluate cohort variant calls. The first data source is the Genome in a Bottle (GIAB) consortium [24,25] which provides a well-characterized set of truth variants. To maximize the diversity of samples and sites used for evaluation, other trios in the cohort were used to compute Mendelian violation rates (with the sole exception being the three person HG002-HG003-HG004 cohort) and the children of GIAB trios were excluded from the GIAB metrics calculations. The second data source is the Clinical Sequencing Evidence-Generating Research (CSER) consortium [26], which contains 929 WGS and 344 WES samples, including 249 WGS trios and 112 WES trios. The third data source is the Population Architecture using Genomics and Epidemiology (PAGE) consortium [27], which contains 313 WGS. The final data source is the recent 30x WGS of 2,504 individuals from 1KGP. We fully withheld the 1KGP cohort from development, only using it as a final independent evaluation set. Within this cohort, a single cryptic trio [28] was used to compute Mendelian violation rates.

We performed analyses both at full sequence coverage as well as in the same cohorts downsampled to 15x coverage. We targeted robust performance across the diversity of sequencing projects by representing cohorts of high and low coverage, WES and WGS, those sequenced by various groups, on various instruments, and across a wide array of ancestries.

### Quality properties of single-sample variant calls

We first investigated variant call properties of 1,248 individuals from the GIAB (n=6), CSER (n=929), and PAGE (n=313) cohorts. The total number of SNPs reported by DeepVariant is lower than GATK4 HaplotypeCaller for nearly all individuals, and the number of indels is also lower for most individuals. However, the Ti:Tv ratio measured in all individuals, and both precision and recall computed separately for SNPs and indels in the six GIAB individuals, are all higher in DeepVariant than GATK4 HaplotypeCaller (**Supplementary Figure 1**).

To illustrate why different single-sample variant callers require separate calibration during joint genotyping, we compared reported Genotype Quality (GQ) scores, defined as the Phred-scaled conditional probability that a genotype is incorrect, to empirical GQ scores derived from the error rate determined from ground truth in the GIAB samples. DeepVariant shows markedly better GQ calibration than GATK4 HaplotypeCaller at all reported GQ scores (**Figure 1A,B**). DeepVariant is well-calibrated both across sequence coverages and when stratified by variant type, with a slight bias toward overconfidence in homozygous alternate SNPs (**Supplementary Figure 2**).

**Figure 1.**
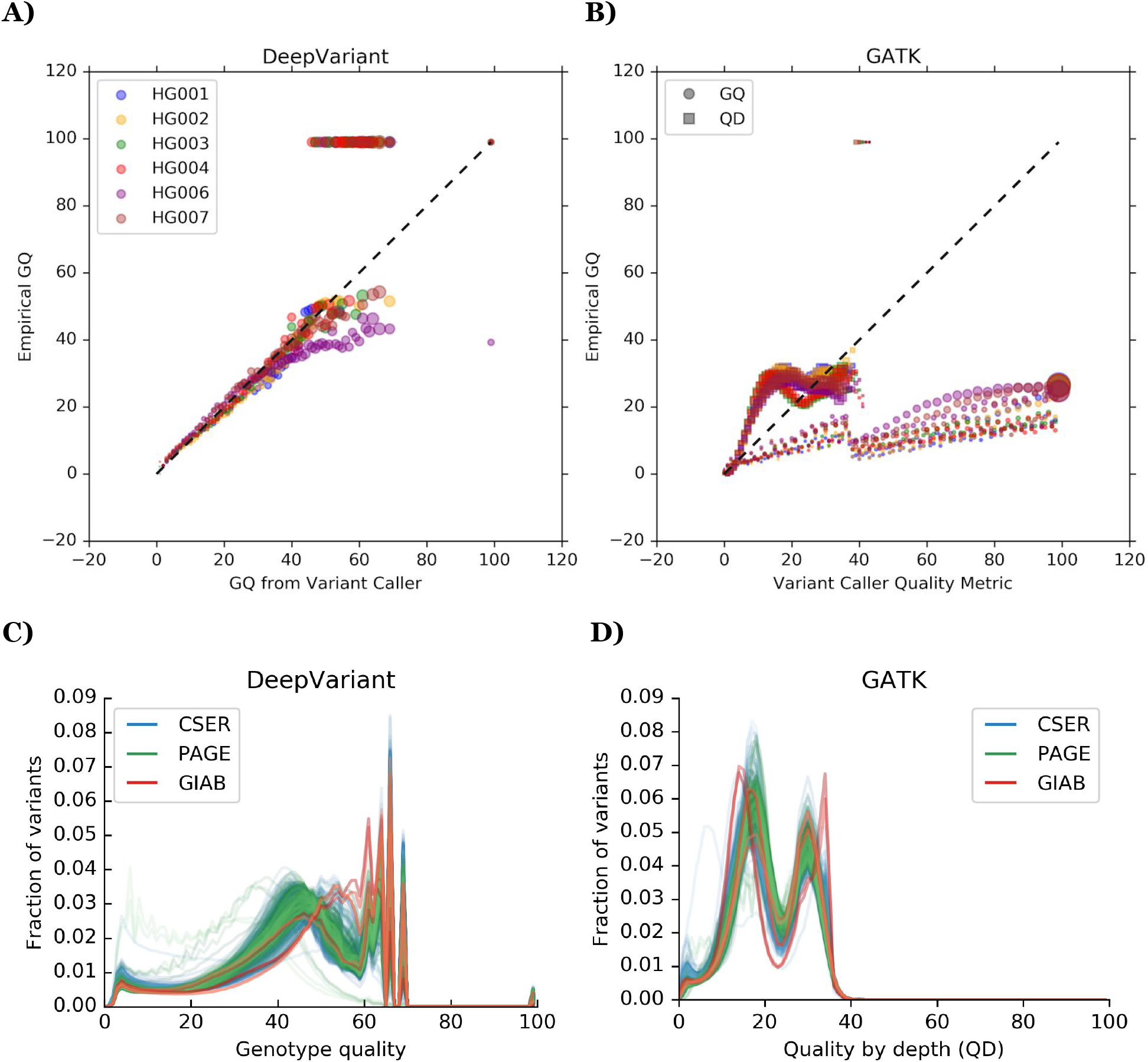
Genotype quality (GQ) distribution properties of PASS variants. **A)** Genotype quality calibration for DeepVariant v0.8.0. Reported GQ is plotted against the empirical GQ calculated using genome-wide GIAB benchmark variant calls at 40x coverage. Each data point is a set of variant calls with the same GQ (x-axis), and the y-axis value is the empirical error rate calculated from the GIAB truth set. Both axes are in Phred-scale. Marker areas are proportional to the square root of the number of variants. The dotted y=x line represents perfect calibration. **B)** Variant calibration for GATK4 HaplotypeCaller. Both reported GQ (circles) and reported QD (variant quality normalized by read depth; squares) are plotted against empirical GQ. Colors correspond to GIAB samples as in A). **C)** Distributions of reported GQ for DeepVariant v0.8.0 in all 1,248 samples computed genome-wide. **D)** Distributions of reported QD for GATK4 HaplotypeCaller in all 1,248 samples computed on chromosome 2 only.

The overall distribution of GQ scores within a sample determines the information content of the field. Substantial variant fractions occur across the GQ spectrum for DeepVariant calls (**Figure 1C**), and the DeepVariant GQ score distribution shifts smoothly toward higher qualities as sequence coverage increases (**Supplementary Figure 3**). In contrast, GATK4 HaplotypeCaller both produces an artifactual ringing behavior for variants with GQ < 99 and frequently reports most variants as GQ=99 (**Supplementary Figure 4**). These results suggest that joint genotyping algorithms may be able to more effectively refine individual genotypes produced by DeepVariant.

The large-scale analysis of genomes and exomes in gnomAD identified variant quality normalized by read depth (QD) as the most important feature for discriminating true variants from artifacts in GATK calls [9]. Consistent with that observation, GATK single-sample QD is less uniformly biased than GQ (**Figure 1B**) and has a more informative distribution across the cohorts (**Figure 1D**).

### Optimized parameters for joint calling

We adapted GLnexus [18] for merging DeepVariant gVCFs because of its computational scalability to large cohorts, access to relevant parameters, performance on allele normalization, and open-source license. To identify optimal parameters for merging DeepVariant gVCFs, we created four custom WGS cohorts of 3, 100, 333, and 1,247 samples at both high coverage (40-50x) and low coverage (15x) on chromosome 2, resulting in eight total cohorts (**Supplementary Table 1**). The cohorts contain five mutually non-descendant GIAB samples used to evaluate benchmark calls and five non-GIAB trios used to compute Mendelian violation rates (**Supplementary Table 1B,C**, **Supplementary Table 2**).

We focused on the four tunable GLnexus parameters (**Supplementary Table 3**) most crucial to optimize: “min_AQ_1_”, the quality threshold applied to each discovered allele; “min_AQ_2_”, the quality threshold applied to alleles whose copy number is at least two; “min_GQ”, the minimum genotype quality to be used for copy number estimates for the alleles; and “revise_genotypes”, a boolean switch indicating whether to use cohort information to re-genotype low quality genotype calls.

We extensively explored parameter configurations using Google Vizier [29] to optimize a multiple metric objective function. Minimization of Mendelian violation rate in trio samples encourages precision genome-wide. Maximization of concordance with GIAB samples, measured as the harmonic mean of SNP and indel F1 scores, encourages both precision and recall in the well-characterized subset of the genome. Together, the joint metric discourages strategies which improve Mendelian violation rate at the expense of genotype errors (for example, by filtering true variant sites). We first performed a Pareto-optimal search using Vizier’s default Bayesian hyperparameter selection algorithm to reduce the problem space, and then explored the reduced space using an exhaustive grid search. Many configurations simultaneously improve both Mendelian violation rate and concordance with GIAB when compared to the GLnexus configuration that performs no variant modification (**Figure 2**, **Supplementary Figure 5**).

**Figure 2.**
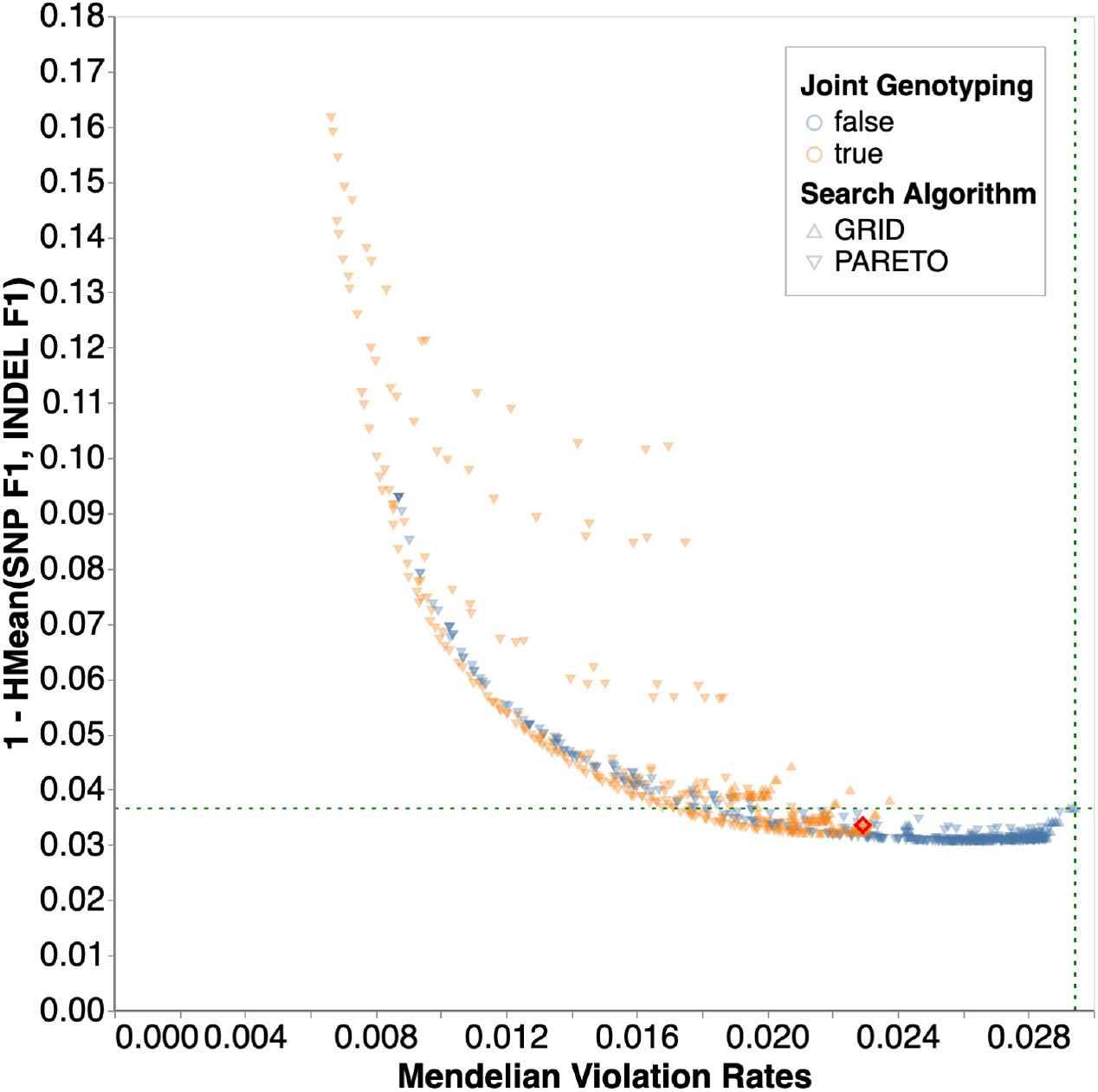
Parameter search for N=1,247, 15x cohort. Each data point represents a unique parameter combination explored by Vizier. The color indicates whether the GLnexus parameter to revise genotypes was true (orange) or false (blue), and the shape represents the search algorithm. The x-axis indicates Mendelian violation rate. The y-axis indicates errors on GIAB through the harmonic mean of SNP F1 and indel F1 (lower is more accurate). Points toward the lower left are more accurate on both metrics. The intersection of the green horizontal and vertical dotted lines indicates the performance using GLnexus with no variant modification (**Supplementary Table 3**). **Supplementary Figure 5** shows the results for all cohort sizes and coverages. The red diamond indicates the parameter set we selected for the optimized DeepVariant+GLnexus pipeline.

The smooth Pareto-optimal boundary (**Figure 2**, **Supplementary Figure 5**) indicates that the tradeoff between recall and precision can be tuned in an application-specific manner. We investigated the extent to which parameter settings are applicable across cohort sizes and sequence coverage by summing the rate of error reduction across five metrics over the “no modification” parameter setting (see **Methods** for details). We selected the best-performing parameter configuration as the “optimized” setting after verifying its strong performance across cohort sizes and sequence coverages (**Supplementary Figures 6-8**).

Next, we compared four variant calling and merging methods across all 8 cohorts. The first and second methods use the GATK4 Best Practices [12,17,20] pipeline and either retain all variants (“GATK-Joint”) or retain only variants that pass variant quality score recalibration (“GATK-VQSR”). The third method uses DeepVariant for single-sample calling and GLnexus to merge the calls, with DeepVariant run with default parameters and GLnexus run in a setting to avoid single-sample variant modification (“DV-GLN-NOMOD”) (**Supplementary Table 3**). The final method uses the optimized version of the DeepVariant+GLnexus pipeline (“DV-GLN-OPT”) (**Supplementary Note 1**). After verifying qualitatively similar callset properties on distinct chromosomes (**Supplementary Figure 9**), we generated all evaluation callsets on chromosome 20 to avoid overlap with the training data from chromosome 2.

The callsets were evaluated on five measures of quality: SNP and indel false discovery rates, false negative rates, and total Mendelian violation rate, for each cohort size and at both 15x and 40x sequence coverage. DV-GLN-OPT equals or exceeds both GATK-based methods in 38 of the 40 measured metrics (**Figure 3**, **Supplementary Table 4**), with only SNP false discovery rates in 15x coverage callsets not uniformly stronger. In the cohort of 1,247 individuals at 40x coverage, DV-GLN-OPT has a 3.0-fold lower Mendelian violation rate (1.7% vs 5.0%), 17.6-fold lower SNP F1 error (0.07% vs 1.23%), and 2.6-fold lower indel F1 error (1.14% vs 2.92%) than GATK-VQSR. DV-GLN-OPT generally, though not strictly, also outperforms DV-GLN-NOMOD.

**Figure 3.**
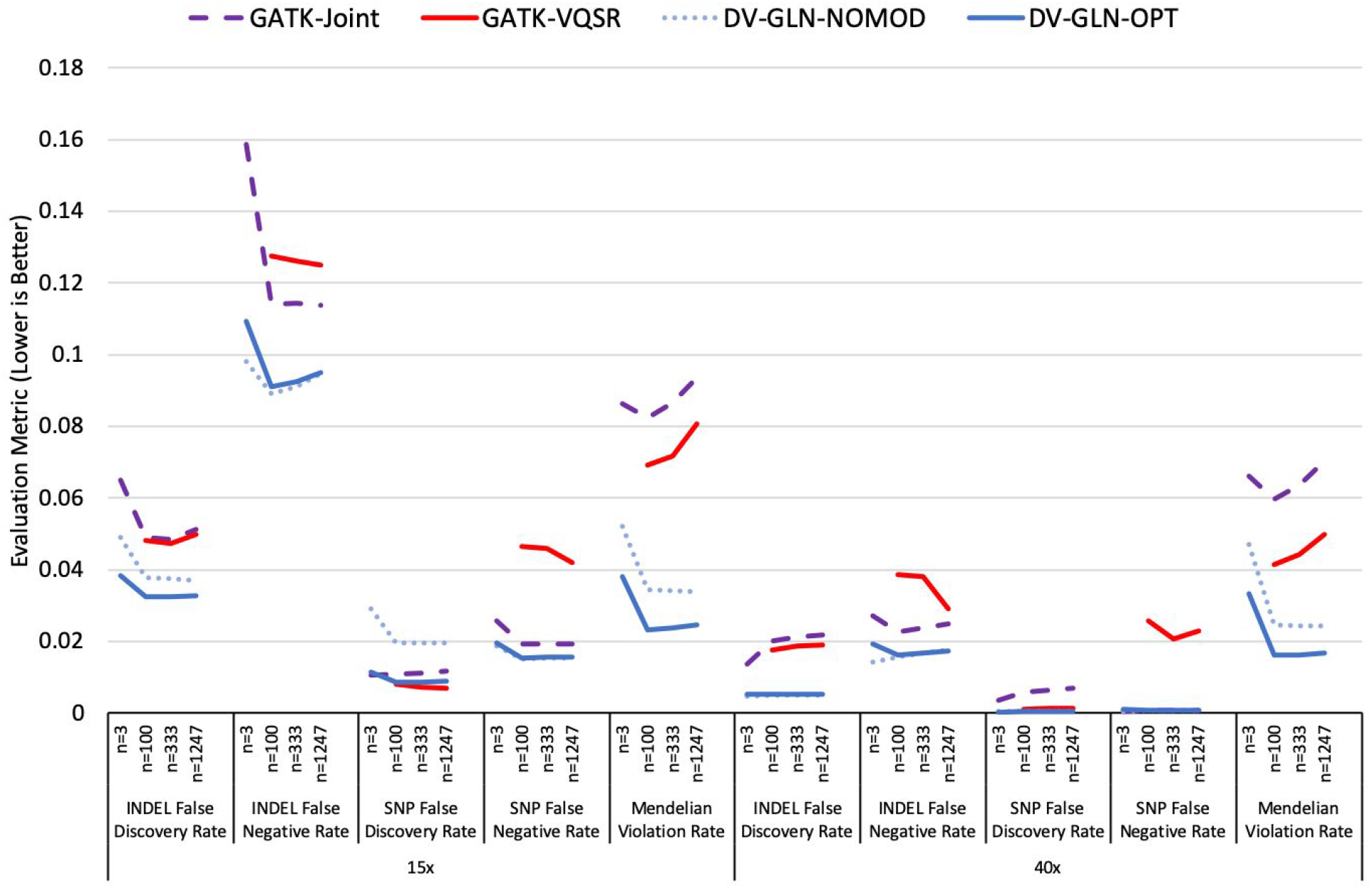
Comparison of four cohort callset creation methods. Four calling and merging pipelines are applied at both 15x and 40x sequence coverage for WGS cohorts of size n=3, 100, 333, and 1247. Five evaluation metrics are presented: Mendelian Violation Rate, SNP False Discovery Rate (1-Precision), SNP False Negative Rate (1-Recall), indel False Discovery Rate, and indel False Negative Rate. In all cases, lower values are better. All evaluation metrics are computed on chr20. See **Supplementary Table 4** for the precise values and the variances of each metric.

We repeated the parameter search technique described above in a single WES cohort of 346 samples (**Supplementary Table 1B**). Similarly to WGS, there exist many configurations that strictly outperform the “no modification” parameter setting (**Supplementary Figure 10**) and the WES-optimized DV+GLnexus pipeline outperforms the GATK4 Best Practices in all metrics (**Supplementary Table 4C**).

### Evaluation on deeply-sequenced 1000 Genomes Project phase 3

To evaluate DV-GLN-OPT in an independent dataset, we generated cohort callsets for high-coverage sequencing reads of the 2,504 samples in 1KGP [21]. We compared the single-sample variant calls from DeepVariant with those of GATK HaplotypeCaller, and the DV-GLN-OPT callset with that of an independently-generated GATK-VQSR pipeline. Each sample has a higher Ti:Tv ratio in the DeepVariant-based variant calls in both cases (**Figure 4A**). Taken together, these results provide indirect evidence that the DeepVariant calls are of higher quality.

**Figure 4.**
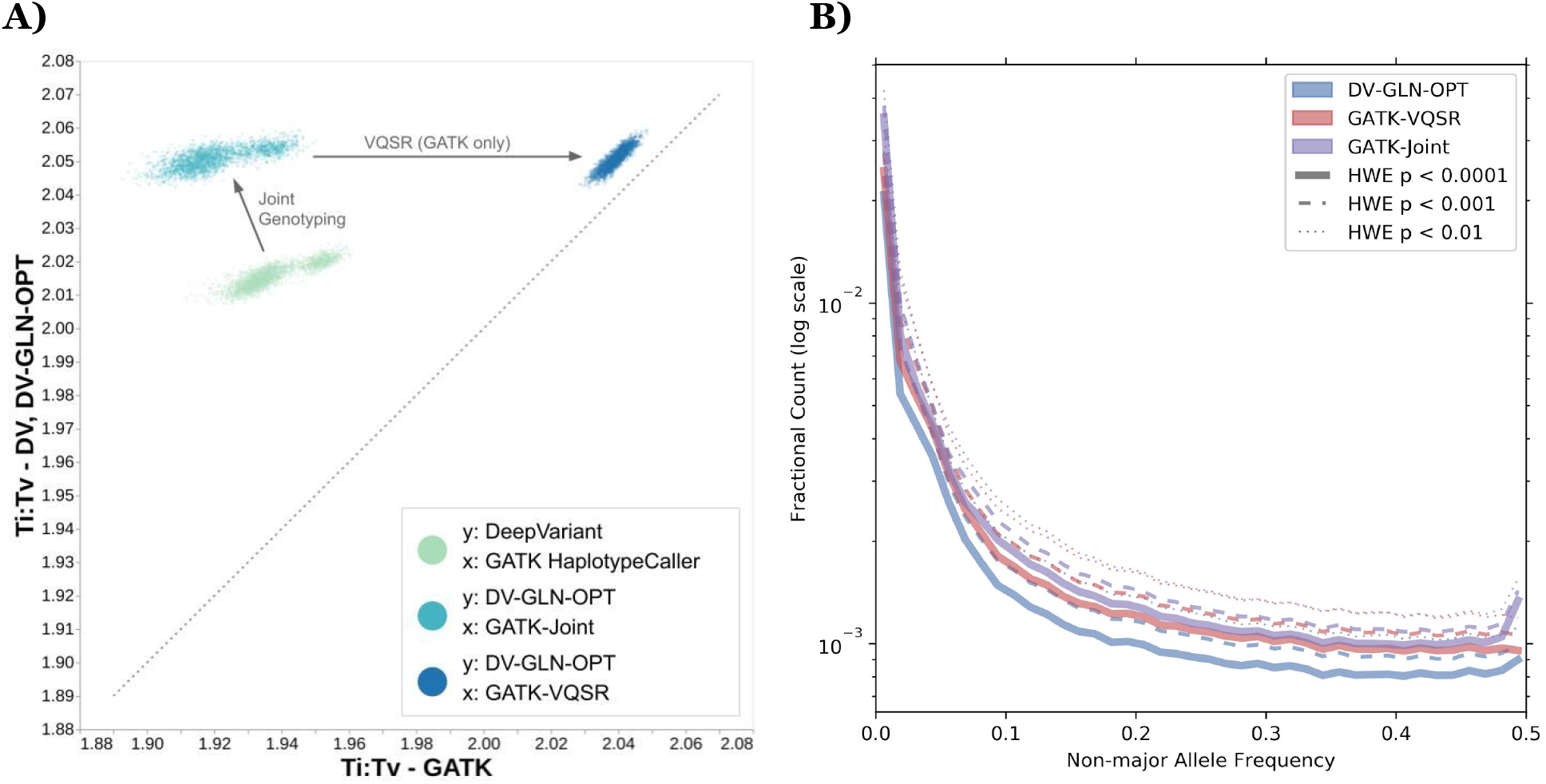
1KGP cohort callset quality. **A)** Ti:Tv ratios of 1KGP samples, from single-sample SNPs and joint-called SNPs, generated by DV-GLN-OPT and GATK pipeline. Each point represents the ratio in one of the 2,504 samples across the whole genome. **B)** Fractional counts of autosomal variants with low HWE p-values, binned by non-major allele frequency in DV-GLN-OPT, GATK-VQSR, and GATK-Joint. The major allele is the allele with the largest allele count in a given variant within the callset. The variants are aggregated in non-major-allele-frequency bins of size 0.0125, and the frequency is clipped at 0.5 for visualization purposes (for all methods the fractional counts in bins after 0.5 is less than 10^-3^).

Overall callset composition depends on the filtering methods applied. The GATK-VQSR callset contains substantially fewer total variants and rare variants compared to both DV-GLN-OPT and GATK-Joint (**Supplementary Figure 11**). To identify recurrent variant calling and genotyping artifacts, we quantified the sites which violate HWE in each callset at various p-value thresholds (**Figure 4B**). Only 6.62% of autosomal sites in the DV-GLN-OPT callset have HWE p<10^-5^ (7,443,684 of 112,451,553 total autosomal sites), compared to 8.05% in GATK-VQSR (8,276,874 of 102,804,074) and 9.77% in GATK-Joint (11,724,367 of 120,046,355). Finally, we observed that the GATK-based callsets limit the maximum number of alleles at any position to six, and thus exclude a number of alleles present at highly variable sites (**Supplementary Figure 11**). Manual inspection confirmed that most highly-multiallelic variants are short tandem repeats of varying lengths, with the bulk of calls attributable to few common alleles but a long tail of additional alleles (**Supplementary Figure 12**).

To further assess 1KGP callset quality, we evaluated Mendelian violation rate within a single cryptic trio present in the cohort [28]. We first verified the trio’s relatedness (NA20882: mother, NA20891: father, NA20900: child, all of Gujarati Indian ancestry). Then, we used this trio to compute Mendelian violations in DV-GLN-OPT, GATK-VQSR, and GATK-Joint (**Figure 5**).

**Figure 5.**
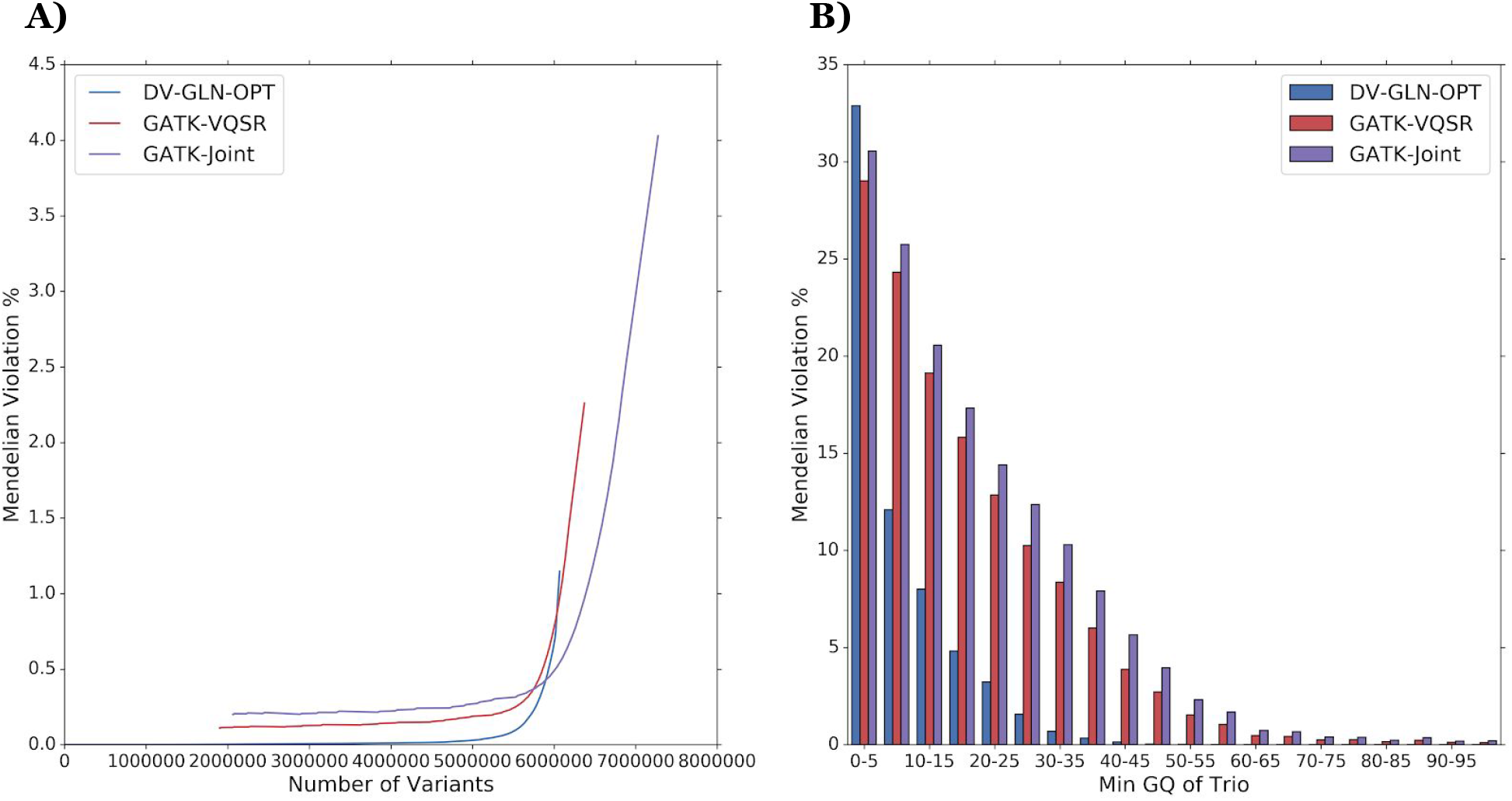
Mendelian violations in autosomes of an inferred trio in 1KGP. **A)** The percentage of variants that violate Mendelian inheritance in the trio NA20900-NA20891-NA20882 as a function of the number of variants considered. Variants are ranked by the minimum GQ within the trio. Callset variants with homozygous reference calls for all three trio samples, and those have indeterminate violation status due to missing genotype calls in the trio, are ignored. **B)** Mendelian violation percentages of the same trio binned by minimum GQ in the trio using bin size 5.

To quantify the improvement to Mendelian violation rate and GQ calibration, we sorted variant calls from most to least confident using the minimum GQ in the trio, independently for each callset. Importantly, variant-level metrics such as QUAL were not used for this analysis because the call qualities for three samples in a trio may differ substantially from a variant quality metric computed over all 2,504 samples. DV-GLN-OPT calls have a lower overall Mendelian violation rate, as evidenced by cumulative Mendelian violation rate plotted as a function of variants retained (**Figure 5A**). While all callsets show decreased Mendelian violation rates as the minimum GQ of the trio is increased (**Figure 5B**), the broader GQ distribution of DeepVariant (**Figure 1**) enables better separation of true and false calls. Remarkably, applying the maximally stringent GQ=99 filter to the GATK-VQSR callset retains only 1.9 million sites (29.8%) at a Mendelian violation rate of 0.11%, whereas the DV-GLN-OPT callset can retain 5.5 million sites (90.6%) at a lower Mendelian violation rate of < 0.1%.

### Evaluation of 1KGP callsets as imputation reference panels

The 1KGP dataset is frequently used for population-based downstream applications, such as genotype phasing and imputation, due to its genetic diversity and large sample size [30–33]. To illustrate how the accuracy of the DV-GLN-OPT callset translates into improved results for these downstream analyses, we assessed the performance of imputing variants using it as a reference panel. We first created reference panels from the deeply sequenced DV-GLN-OPT and GATK-VQSR 1KGP callsets described in the previous section, by applying identical minimal transformations to both cohort VCFs (see **Methods**) and phasing the callsets with Eagle2 [34].

The DV-GLN-OPT panel contains 4.69% more variant sites than the GATK-VQSR panel generated from the same source. More than 99% of the GATK-VQSR panel sites are present in the DV-GLN-OPT panel, while fewer than 95% of the DV-GLN-OPT panel sites are present in the GATK-VQSR panel (**Figure 6A**).

**Figure 6.**
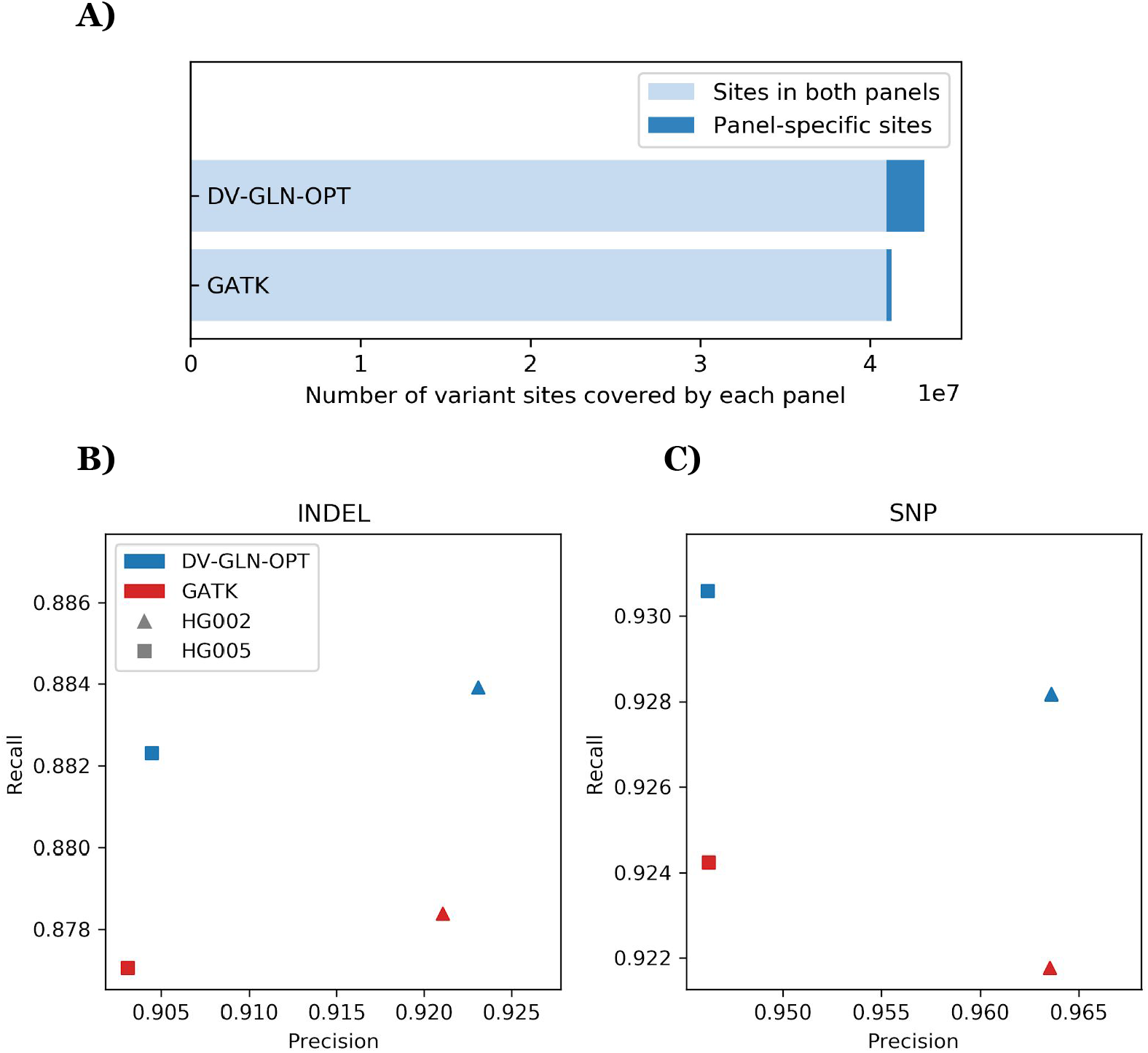
Imputation accuracy of 1KGP reference panel. **A) Variant sites covered by DV-GLN-OPT and GATK panel.** The DV-GLN-OPT reference panel generated from 1KGP samples covers 43,181,562 variant sites, while the GATK panel from the same samples covers 41,247,330 sites. The intersection of the two panel regions (marked in light blue) covers 40,972,007 sites, which is 94.88% of the DV-GLN-OPT panel and 99.33% of the GATK panel. **B) Imputed genotype accuracy for indels.** The accuracy of the imputed variants are measured by computing concordance with the GIAB benchmark calls using hap.py. Blue colored markers are from DV-GLN-OPT panel while the red markers are from GATK panel. The shaped markers show precision and recall computed across the GIAB evaluation region for two samples. **C) Imputed genotype accuracy for SNPs.** Shapes and colors as in B).

To evaluate the imputation quality of the two reference panels, we extracted variant calls for the ~710k sites assayed by the Illumina Infìnium OmniExpress-24 microarray for two GIAB child samples (HG002 and HG005) in their benchmark regions. For each of the DV-GLN-OPT and GATK-VQSR reference panels, we phased the pseudo-microarray variants with Eagle2 and imputed the phased variants into the panel with Beagle 5.0 [35].

The imputed variant calls were scored in two evaluation regions (**Supplementary Table 5**). The first evaluation region, hereafter the “GIAB evaluation region,” comprises the GIAB benchmark regions common to both HG002 and HG005, agnostic to either reference panel. This measures both the accuracy of the imputed genotypes and the number of benchmark variants absent in the reference panel. The second evaluation region, hereafter the “panel evaluation region,” comprises a subset of the GIAB evaluation region additionally present in both the DV-GLN-OPT and the GATK-VQSR reference panels. This allows a direct comparison of variants, but provides limited information about overall individual panel quality.

The DV-GLN-OPT panel outperforms the GATK panel in F1 score in all eight experiments (two samples, two variant types, and two evaluation regions). Of note, DV-GLN-OPT produces substantially higher recall than GATK-VQSR for both indels and SNPs when evaluated in the GIAB evaluation region (**Figure 6B,C**). The DV-GLN-OPT panel produces on average 4.41% fewer false negative indels and 8.28% fewer false negative SNPs, while maintaining superior indel precision and indistinguishable SNP precision. As expected, evaluation metric differences are more subtle in the panel evaluation region, but the DV-GLN-OPT panel produces higher F1 scores for both samples and for both indels and SNPs (**Supplementary Table 5**).

### Cost benchmarking

In large-scale sequencing projects, the temporal and financial cost of running bioinformatics tools can be prohibitively large. To compare the computational cost of the DeepVariant-GLnexus and GATK pipelines, we reanalyzed chromosome 22 in the 2,504 1KGP samples. Starting from the aligned sequencing reads, we ran DeepVariant and GATK HaplotypeCaller to produce gVCFs on a separate virtual machine with a fixed machine type for each sample, using the Docker images published by the respective tool developers.

Using machines with 8 virtual CPUs (vCPUs) each, DeepVariant finished each chr22 sample using 40% less average time elapsed (20.0 minutes) without GPU/TPU acceleration, than GATK HaplotypeCaller (33.2 minutes) (**Figure 7A**). The difference is mainly attributable to DeepVariant’s efficient internal multithreading (**Figure 7B**, **Supplementary Table 6**). This implies that one can easily assign more vCPUs to each cloud machine to get a speedup almost proportional to the increased resources (**Supplementary Figure 13**), without requiring an external workflow that splits the chromosome into smaller shards. Note that the cost difference between the two callers would expand significantly if the Base Quality Score Recalibration (BQSR) preprocessing step were included for GATK, which is part of GATK Best Practices but not recommended for DeepVariant.

**Figure 7.**
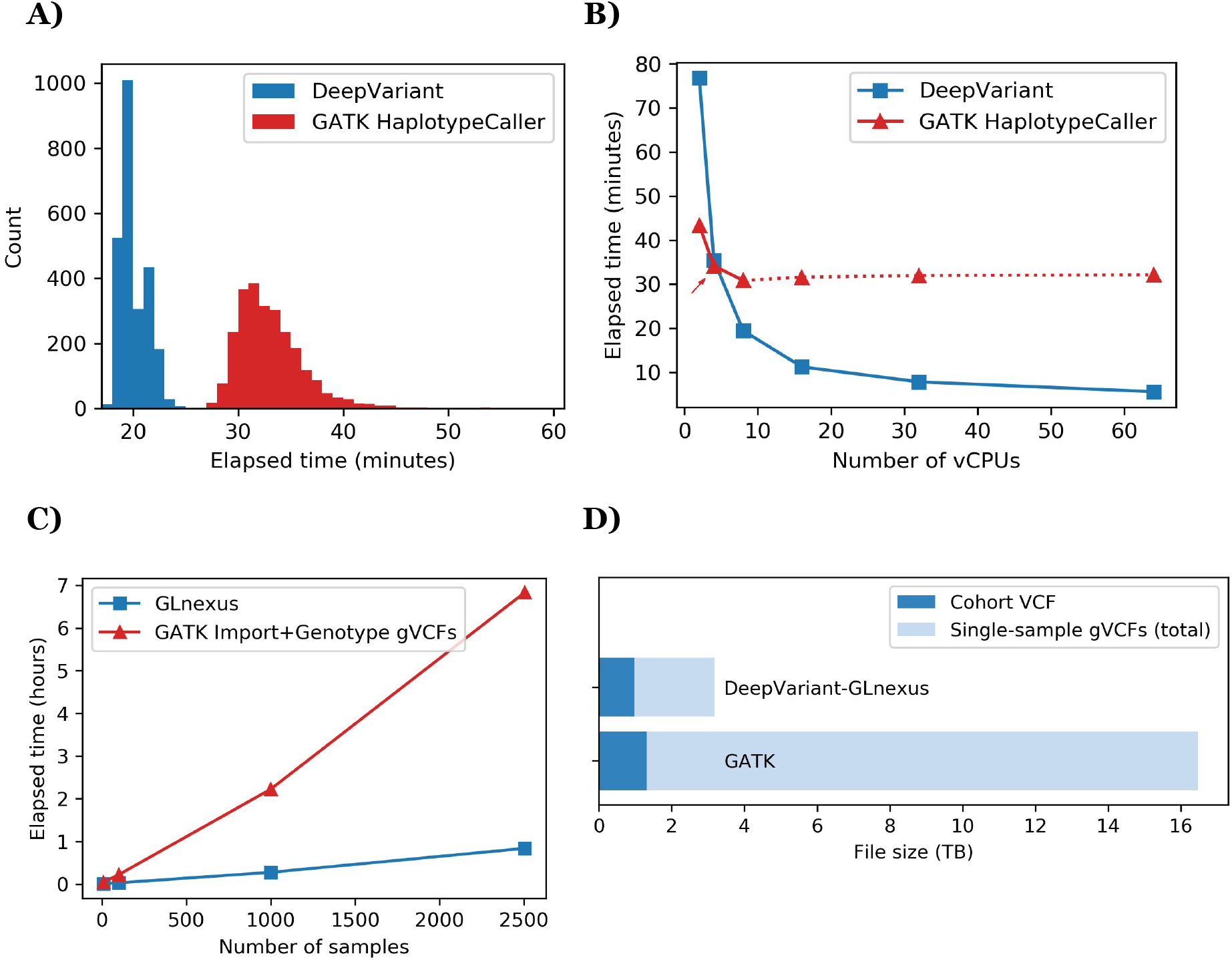
Cost benchmarking DeepVariant-GLnexus and GATK pipeline. **A)** Distribution of elapsed real times to generate single-sample gVCF (chr22 only) from aligned reads across N=2,504 1KGP samples, using DeepVariant and GATK HaplotypeCaller (BQSR not included) in 8-vCPU machine. GPU/TPU acceleration was *not* used for DeepVariant. **B)** Elapsed real times to generate gVCF (chr22 only) of one sample (NA12878) using a cloud machine with varying number of vCPUs, with DeepVariant and GATK HaplotypeCaller (excluding BQSR). The default value for HaplotypeCaller’s HMM multithreading flag (`--native-pair-hmm-threads`) is 4 (red arrow) and it was practically ineffectual for 16 vCPUs and more (red dotted lines). **C)** Elapsed real times to merge the chr22 gVCF files from (A) into a cohort VCF for N={10, 100, 1000, 2504} nested subsets of the 1KGP samples, using GLnexus (for DeepVariant gVCFs) and GATK GenomicsDBImport + GenotypeGVCFs (for HaplotypeCaller gVCFs). GATK VQSR step was not included. **D)** The file sizes of the whole-genome cohort VCFs and the single-sample gVCFs of 1KGP samples from DeepVariant-GLnexus and GATK pipeline.

Next, we processed the N=2,504 sample chromosome 22 gVCF files to produce cohort VCF files using GLnexus (from DeepVariant gVCFs), on the one hand, and GATK’s GenomicsDBImport and GenotypeGVCFs tools (from HaplotypeCaller gVCFs), on the other. While GLnexus supports internal multithreading, the two GATK tools are effectively single-threaded and require an external parallelization workflow to achieve practical runtimes, which we reproduced based on the developers’ specifications (subdividing the length of chromosome 22 to scatter across processes). Still, using a single 32-vCPU virtual machine, GLnexus is 8 times faster (0.84 hours) than the equivalent GATK tools (6.83 hours), with superior scalability to larger cohorts (**Figure 7C**). For this benchmark we did not run Variant Quality Score Recalibration (VQSR) for GATK, which is a recommended step after GenotypeGVCFs in its Best Practices and will add additional runtime to its pipeline.

Another relevant cost to users of these pipelines is the cost of storing the artifacts from them. In the standard block-compressed variant call format [36,37], the total size of the DeepVariant gVCFs for all chromosomes of 1KGP samples is 7 times smaller (total 2.20TB, average 878MB/sample) than GATK HaplotypeCaller gVCFs (total 15.16TB, average 6,053MB/sample) (**Figure 7D**), which is a result of DeepVariant’s efficient quantization of the reference records. Moreover, the final cohort VCF from DeepVariant-GLnexus pipeline is also 26% smaller (0.97TB) than the one from GATK pipeline (1.32TB). This reduction in file sizes directly translates to a similar ratio of cost savings in cloud storage services. To further reduce the sizes of the cohort VCF, one may consider using the BCF (binary VCF) format or other data formats designed for a large number of samples [38–43].

## Discussion

Population-scale sequenced cohorts are foundational resources for many genetic analyses, including genotype-phenotype discovery, variant interpretation, and genotype imputation. As sequencing projects have grown to include hundreds of thousands of samples, the need for highly accurate variant calls and computationally efficient merging algorithms is increasingly acute. By optimizing GLnexus [18] to merge single-sample DeepVariant calls, we demonstrated that the superior accuracy [15] and generalizability across sequencing methods [44] of DeepVariant can generate more accurate cohort callsets at large scale at lower cost. The callset quality metrics of the optimized pipeline consistently outperformed the GATK Best Practices across a range of cohort sizes and sequence coverages. In addition, we showed that variant confidences are well calibrated to Mendelian violation rate, allowing tuning of callsets for very high precision or for high recall.

When optimizing callset creation, we investigated callset quality stratified by both sequencing coverage and cohort size. Results within a given sequencing coverage were qualitatively similar regardless of cohort size, suggesting that a major driver of parameter sensitivity is the distribution of individual call confidence estimates. Even so, when optimizing equally between benchmark callset accuracy and Mendelian violation rate, we observed a single parameter set that provides strong performance across the range of WGS cohorts analyzed.

Although we demonstrated the strength of the DeepVariant+GLnexus method, there are multiple areas for future improvement. At both 15x and 40x coverage, the precision-recall curves of SNPs vs. indels is markedly different. As expected, parameter variation can tune SNPs to improve recall at the expense of precision or vice versa. In contrast, indels appear to have nearly globally optimal parameters, suggesting that distinct handling of the two variant classes may further improve callset quality. Additionally, the tunable GLnexus parameters affect allele harmonization and genotyping, but do not apply any hard filtering to output calls. We observed an overrepresentation of Mendelian violations at very low GQ values, indicating that direct omission of low quality sites or genotypes also may improve callset quality. Finally, a small fraction (0.4%) of autosomal sites contain seven or more total alleles, and typically represent short tandem repeats [45]. While these sites likely capture some of the known hypervariability of these regions, this benefit is weighed against the practical difficulty of representing and analyzing these sites in downstream applications.

We used the recent deep sequencing of 1KGP to perform an orthogonal analysis on a publicly available dataset. The resulting optimized DeepVariant+GLnexus callset possesses superior metrics to a GATK Best Practices callset generated independently, including a 32% reduction in sites violating HWE at p<10^-5^. Moreover, an imputation reference panel derived from the DeepVariant+GLnexus callset results in higher imputation accuracy, which shows that improving cohort-level variant calls yields improved performance in downstream applications.

Both the cohort callset and all DeepVariant single-sample calls are freely available at gs://brain-genomics-public/research/cohort/1KGP/. To our knowledge, this is the most accurate 1KGP callset currently available, and as such has substantial utility within the genomics community for studies of genetic variation. Furthermore, we hope this resource spurs additional innovation in the development and evaluation of population-scale cohorts.

## Methods

### Data acquisition and preparation

#### Reference genome

Throughout this study we used the human GRCh38 reference genome that contains the “no alt” analysis set and human decoy sequences from hs38d1 (GCA_000786075.2) (ftp://ftp.ncbi.nlm.nih.gov/genomes/all/GCA/000/001/405/GCA_000001405.15_GRCh38/se_qs_for_alignment_pipelines.ucsc_ids/GCA_000001405.15_GRCh38_no_alt_plus_hs38d1_an_alysis_set.fna.gz; ftp://ftp.ncbi.nlm.nih.gov/genomes/all/GCA/000/001/405/GCA_000001405.15_GRCh38/seqs_for_alignment_pipelines.ucsc_ids/README_analysis_sets.txt).

#### Genome in a Bottle

We used sequencing reads and benchmark callsets of the seven individuals currently in GIAB. HG001 (a female of Utah/European ancestry) and HG002 (Ashkenazi Jewish son) samples were sequenced by Illumina HiSeq 2500 in Rapid Mode (v1) with 2×148bp read length and ~50x coverage, and were aligned to GRCh38 by BWA-MEM version 0.7.17 [46]. The Ashkenazi Jewish parent samples, HG003 and HG004, were sequenced with Illumina HiSeq 2500 in Rapid mode (v2) with 2×250 paired-end reads and ~40-50x coverage, and were aligned to GRCh38 using Novoalign version 3.02.07 (www.novocraft.com/products/novoalign). HG005, the Chinese son sample, was used only for imputation evaluation based on the benchmark callset. The Chinese parent samples HG006 and HG007 were sequenced by Illumina HiSeq 2500 in Rapid mode (v1) with 2×148 paired end reads to 100x coverage, mapped to GRCh38 with BWA-MEM, and downsampled to ~40x coverage to match the coverage of other samples using samtools v1.6 [37].

#### Clinical Sequencing Evidence-Generating Research

We downloaded SRA files for 929 WGS samples (among 931 WGS samples, SRR6706856 and SRR4370311 repeatedly failed to download) and 346 WES samples from the CSER project from dbGaP (project ID: 20844). All samples were sequenced with the Illumina HiSeq X platform. We generated FASTQ files using the “fastq-dump” command in the NCBI SRA Toolkit (github.com/ncbi/sra-tools). Finally, all FASTQ files were mapped to GRCh38 with BWA-MEM version 0.7.17 and duplicate reads were marked by samblaster version 0.1.24 [47].

We identified all mother-father-child trio samples in the CSER dataset using the “Library_Name” field in the associated SRA Run Table. The values of the field are formatted as “A-[Family ID]-[Family relation]”. Of 249 disjoint trios we identified, we randomly selected five trios (15 total individuals, IDs in **Supplementary Table 2**) among all non-outlier trios to use for Mendelian violation rate estimation during callset evaluation. To define non-outlier samples, we examined six variant summary statistics for each sample: the number of records, the number of SNPs, the number of indels, the Ti:Tv ratio, the mean SNP quality, and the mean indel quality. Non-outliers are defined as the samples for which all six statistics are within one standard deviation of the mean (i.e. the magnitude of the Z-score is at most one).

#### Population Architecture Using Genomics and Epidemiology

We processed the PAGE samples in the same way as those from CSER. We downloaded 313 sample SRA files from dbGaP (project ID: 17123), which were also generated by Illumina HiSeq X platform, converted them to FASTQ, mapped them to GRCh38, and marked duplicates as described above. We also generated a subset we call *PAGE80* by randomly selecting 80 non-outlier samples among the 313 samples (**Supplementary Table 7**). The same six summary statistics were used to select non-outliers, except that we used a maximum Z-score magnitude of 1.25 (instead of one) to include more samples.

Using the 6 GIAB samples, 929 CSER samples, and 313 PAGE samples above, we created custom cohorts of size 3, 100, 333, and 1247 (**Supplementary Table 1**) for which both GIAB concordance and Mendelian violation rate could be evaluated. We used the GIAB benchmark variant version 3.3.2 for GRCh38 to evaluate concordance. Finally, we created 15x autosomal coverage BAMs from all BAM files from GIAB, CSER, and PAGE datasets by downsampling full BAMs with samtools v1.6 using the “samtools view -s” command.

#### 1000 Genomes Project

The 2,504-sample cohort callset we release is based on the recent deep sequencing of the 1KGP phase3 samples by New York Genome Center. The input reads were sequenced at 30x coverage using the new Illumina NovaSeq 6000 system with 2×150bp reads, and then aligned to GRCh38 using BWA-MEM v0.7.15 [46]. More details about their pipeline can be found on EBI 1000 Genomes FTP (ftp://ftp.1000genomes.ebi.ac.uk/vol1/ftp/data_collections/1000G_2504_high_coverage/201_90405_NYGC_b38_pipeline_description.pdf). We used samtools v1.6 to convert the original CRAM files to BAM format for downstream tasks.

### DeepVariant

We used DeepVariant v0.8.0 and the publicly-released WGS model v0.8.0 to generate the single-sample variant calls for all samples in GIAB, CSER, and PAGE. A single-line command to run DeepVariant in a pre-built docker container is available on the DeepVariant public repository (github.com/google/deepvariant). The DeepVariant calls for the sample from 1000 Genomes Project were generated using a custom model trained exclusively for the NovaSeq platform. Both the custom model and all single-sample DeepVariant calls generated by it are publicly available, as described in **Data availability**.

### GLnexus

To merge and evaluate the multiple cohorts in parallel, we deployed the open-source GLnexus algorithm using Apache Beam (beam.apache.org) on Google internal compute clusters. The Beam-based pipeline abstracts away the need to specify multi-threading on a single machine (as is done in the open-source GLnexus), and deploys hundreds of different parameter configurations on thousands of CPUs. The pipeline produces identical scientific results to the open-source GLnexus v1.2.2 when run with the same parameters. We used chromosome 2 to optimize the pipeline, and computed final performance benchmarks separately on chromosome 20. The optimized DeepVariant parameters from this study are included in open-source GLnexus v1.2.2 or later versions in two presets: `--config DeepVariantWGS` for WGS and `--config DeepVariantWES` for WES. After installing the GLnexus command line tool, users can merge DeepVariant calls in these optimized setups using a single command like

~~~
$ glnexus_cli --config DeepVariantWGS deepvariant.*.g.vcf.gz > cohort.bcf
~~~

### GATK

We followed GATK Best Practices version 4.1.2.0 to establish the baseline performance of each callset generated from GIAB, CSER, and PAGE data. Starting from the BAM files, prepared as described above, we ran HaplotypeCaller in GVCF mode to call single-sample variants, followed by GenomicsDBImport to consolidate the GVCF files, GenotypeGVCFs to jointly genotype the cohort, and finally VariantRecalibrator and ApplyVQSR for variant quality score recalibration (VQSR). The full details of the VQSR parameters can be found in **Supplementary Note 2**. We performed the steps on each chromosome separately in parallel and combine calls at the end to speed up the process. The final performance benchmarks were performed on chromosome 20.

For the 1KGP samples, we downloaded the released GATK cohort callset (ftp://ftp.1000genomes.ebi.ac.uk/vol1/ftp/data_collections/1000G_2504_high_coverage/working/20190425_NYGC_GATK/). This was generated by the New York Genome Center using samtools v1.3.1, Picard v2.4.1, and GATK v3.5 (BaseRecalibrator, HaplotypeCaller, GenotypeGVCFs, VariantRecalibrator, and ApplyRecalibration). The complete description of the pipeline used to generate this callset is available on EBI 1000 Genomes FTP (ftp://ftp.1000genomes.ebi.ac.uk/vol1/ftp/data_collections/1000G_2504_high_coverage/working/20190425_NYGC_GATK/1000G_README_2019April10_NYGCjointcalls.pdf).

### Single-sample variant call statistics

We used bcftools v1.9 (samtools.github.io/bcftools) and hap.py v0.3.9 (github.com/illumina/hap.py) to generate basic call statistics from single-sample VCFs in the GIAB, PAGE, and CSER datasets.

### GLnexus parameter optimization

We used Google Vizier [29], a Google-internal service for performing black-box optimization, for optimizing the configurable parameters of GLnexus (**Supplementary Table 3**).

The first iteration of parameter search used the *Pareto-optimal* search algorithm, where we set two optimization objectives: maximizing GIAB benchmark call concordance and minimizing the rate of Mendelian violations. GIAB benchmark call concordance was defined as the harmonic mean of the SNP F_1_ score (which in turn is defined as the harmonic mean of SNP recall and precision values) and the indel F_1_ score. The precision/recall of the two types of variants was defined as the arithmetic mean of the precision/recall values for the GIAB benchmark samples (three samples for the cohorts of size three, and five samples for all other cohorts). The purpose of the first iteration of parameter search was to explore the general trends and reduce the search space volume. To this end, we explored a wide range of possible parameter values: “min_AQ_2_” could be any integer from 0 to 50 (inclusive), “min_AQ_1_” could be any integer from “min_AQ_2_” to 80 where the difference between “min_AQ_1_” and “min_AQ_2_” was at most 30, “min_GQ” was a multiple of 10 from 0 to 50 (because GLnexus quantizes this value as a multiple of ten), and “revise_genotypes” could be True or False.

After manually investigating points on the Pareto optimal frontier of the above search we substantially reduced the search space as follows: “min_AQ_2_” between 0 and 20, “min_AQ_1_” defined by (0 ≤ min_AQ_1_ - min_AQ_2_ ≤ 20), “min_GQ” in {0, 10, 20}, and “revise_genotypes” could be True or False. In this reduced search space, we performed exhaustive grid search where the size of the grid for “min_AQ_1_” and “min_AQ_2_” was 5 (so the values are multiples of 5). This resulted in 150 (=5×5×3×2) total configurations. We merged all eight cohorts with all possible configurations in this space, resulting in 1,200 total experiments.

Formally, we have the following five evaluation metrics, each of which is a function of the read coverage, cohort size, and configuration parameters:

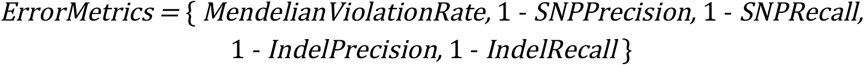

The value of every metric is between 0 to 1, where 0 is the most desirable. Note that 1-precision is also called the *false discovery rate* and 1-recall is called the *false negative rate*.

Let *P* be the set of all parameter tuples within our search space, namely,

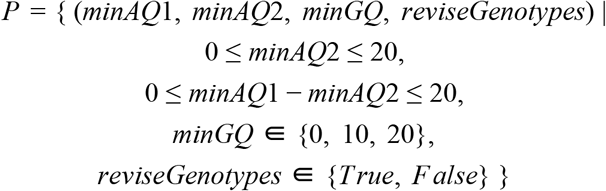

For each parameter set *p* in *P*, we define the objective function *L*(*p*) by this formula:

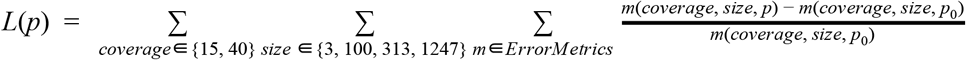

where *p_o_* is the GLnexus parameter with no modification of input calls, namely *p_0_*=(0, 0, 0, *False*). Finally we search for the parameter *p\**that minimizes the objective.

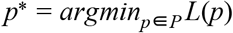

This implies that we try to maximize the *rate of error reduction* per metric over the “no modification” parameter setting, and sum them with equal weights.

The optimized DeepVariant+GLnexus callsets use this *p\** configuration (see **Supplementary Note 1** for the parameter values).

### 1KGP imputation reference panel creation

We generated the 1KGP reference panels from DV-GLN-OPT and GATK-VQSR callsets by applying identical minimal transformations to them and phasing them with Eagle[34]. We followed a standard pipeline for generating a reference panel recommended in Eagle’s website (data.broadinstitute.org/alkesgroup/Eagle/#x1-300005.3). Starting from each cohort callset, we removed singleton variants and kept only variants with either “PASS” or “.” (empty) filter. We also converted multi-allelic variants to multiple biallelic variants and removed duplicate variants, as required by Eagle, using bcftools v1.9 (samtools.github.io/bcftools). Finally, we ran Eagle v2.4.1 with the hg38 genetic map file released with the software, without supplying any additional reference panel. A script for running all the above steps can be found in **Supplementary Note 3**.

### Imputing pseudo-microarray variant calls

To evaluate 1KGP imputation reference panels, we first generated pseudo-microarray variant calls from GIAB benchmark variants of HG002 and HG005 in the benchmark regions by extracting the variant sites used by a popular commercial microarray kit *(Illumina Infnium OmniExpress-24).* We obtained the microarray sites from the CSV manifest file (GRCh38 version) on Illumina’s official website (ftp://webdata2:webdata2@ussd-ftp.illumina.com/downloads/productfiles/humanomniexpress-24/v1-2/infinium-omniexpress-24-v1-2-manifest-file-csv.zip) and converted it to a VCF format using Illumina’s *GTCtoVCF* tool (github.com/Illumina/GTCtoVCF).

Starting from GIAB v3.3.2 benchmark variants for HG002 and HG005, we removed all existing phasing information (genotypes marked with the pipe symbol or the PS format field) from the VCF, extracted the high-confidence variants in the microarray sites using bcftools v1.9, and added homozygous-reference genotypes to all microarray sites not present in GIAB VCFs. We phased the resulting pseudo-microarray variants with Eagle v2.4.1 using the reference panel to evaluate (DV-GLN-OPT or GATK-VQSR) and the hg38 genetic map file released with Eagle.

Finally, we imputed the phased pseudo-microarray variants with *Beagle* v5.0 [35] using the same reference panel used in the phasing step. A complete script for running Beagle can be found in **Supplementary Note 4**.

## Declarations

### Ethics approval and consent to participate

Not applicable.

### Consent for publication

Not applicable.

### Availability of data and materials

The optimized DeepVariant merging parameters from this study are included in open-source GLnexus v1.2.2 or later versions in two presets: `--config DeepVariantWGS` for WGS and `--config DeepVariantWES` for WES. After installing the GLnexus command line tool, users can merge DeepVariant calls in these optimized setups using a single command such as:

~~~
$ glnexus_cli --config DeepVariantWGS deepvariant.*.g.vcf.gz > cohort.bcf
~~~

The DV-GLN-OPT callset and the DeepVariant single-sample calls for the 1KGP cohort are publicly available in Google Cloud Storage in the gs://brain-genomics-public/research/cohort/1KGP/ folder (console.cloud.google.com/storage/browser/brain-genomics-public/research/cohort/1KGP). Instructions for accessing this public data can be found in Google Cloud Storage documentation (cloud.google.com/storage/docs/access-public-data).

### Author contributions

C.Y.M. designed the study. H.L. and T.Y. implemented the code. T.Y., H.L., M.F.L., and C.Y.M. performed experiments. T.Y., H.L., P-C. C., M.F.L., A.C., and C.Y.M. analyzed the results. T.Y., A.C., M.F.L., and C.Y.M. wrote the manuscript with contributions from all authors.

## Acknowledgements

We thank Babak Alipanahi, Thomas Colthurst, Greg Corrado, and Mark DePristo for helpful discussions and feedback, and the New York Genome Center and its sponsors for the deep resequencing of the 1KGP samples.

## PAGE dataset acknowledgment

Samples and data of The Charles Bronfman Institute for Personalized Medicine (IPM) BioMe BioBank used in this study were provided by The Charles Bronfman Institute for Personalized Medicine at the Icahn School of Medicine at Mount Sinai (New York). Phenotype data collection was supported by The Andrea and Charles Bronfman Philanthropies. Funding support for genotyping, which was performed at The Center for Inherited Disease Research (CIDR), was provided by the NIH (U01HG007417). The datasets used for the analyses described in this manuscript were obtained from dbGaP at http://www.ncbi.nlm.nih.gov/gap through dbGaP accession number phs000925.v1.p1.

## CSER dataset acknowledgment

These results are in whole or part based upon data generated by the Clinical Sequencing Exploratory Research (CSER) consortium established by the NHGRI. Funding support was provided through cooperative agreements with the NHGRI and NCI through grant numbers U01 HG007301(Genomic Diagnosis in Children with Developmental Delay). Information about CSER and the investigators and institutions who comprise the CSER consortium can be found at http://www.genome.gov/27846194.

