## Supplementary Material for "Accurate, scalable cohort variant calls using DeepVariant and GLnexus"

Supplementary Figures **1**

Supplementary Tables **14**

Supplementary Notes **24**

Supplementary Note 1: "DV-GLN-OPT" optimized GLnexus WGS configuration 24
Supplementary Note 2: GATK VQSR parameters 26
Supplementary Note 3: Reference panel creation 28
Supplementary Note 4: Genotype imputation 29
Supplementary Note 5: Software versions 29

### Supplementary Figures

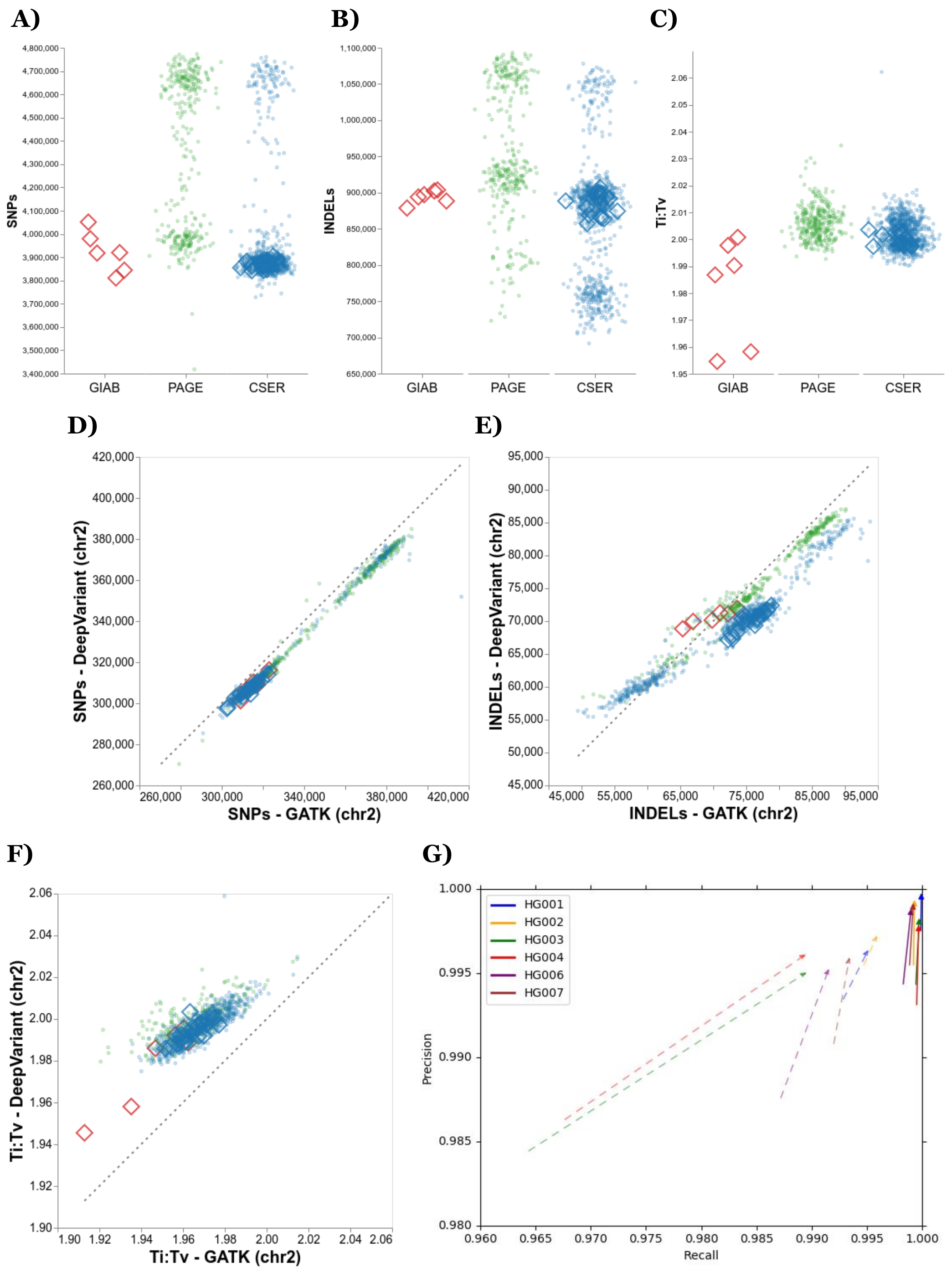


**Supplementary Figure 1. Single-sample call statistics for GIAB, PAGE, and CSER datasets. A, B, C)** The number of SNPs (A), indels (B), and Ti:Tv ratio (C) reported in each individual genome-wide using DeepVariant. Diamond markers indicate samples used for evaluation (GIAB samples for benchmark call accuracy and CSER samples for Mendelian violation rate). **D, E, F)** Comparison of DeepVariant and GATK4 HaplotypeCaller single-sample calls for number of SNPs (D), indels (E), and Ti:Tv ratio (F) computed on chromosome 2. **G)** Comparison of GATK4 HaplotypeCaller (line starts) and DeepVariant (arrowheads) recall and precision scores for SNPs (solid lines) and indels (dashed lines) computed in the GIAB samples on chromosome 2.

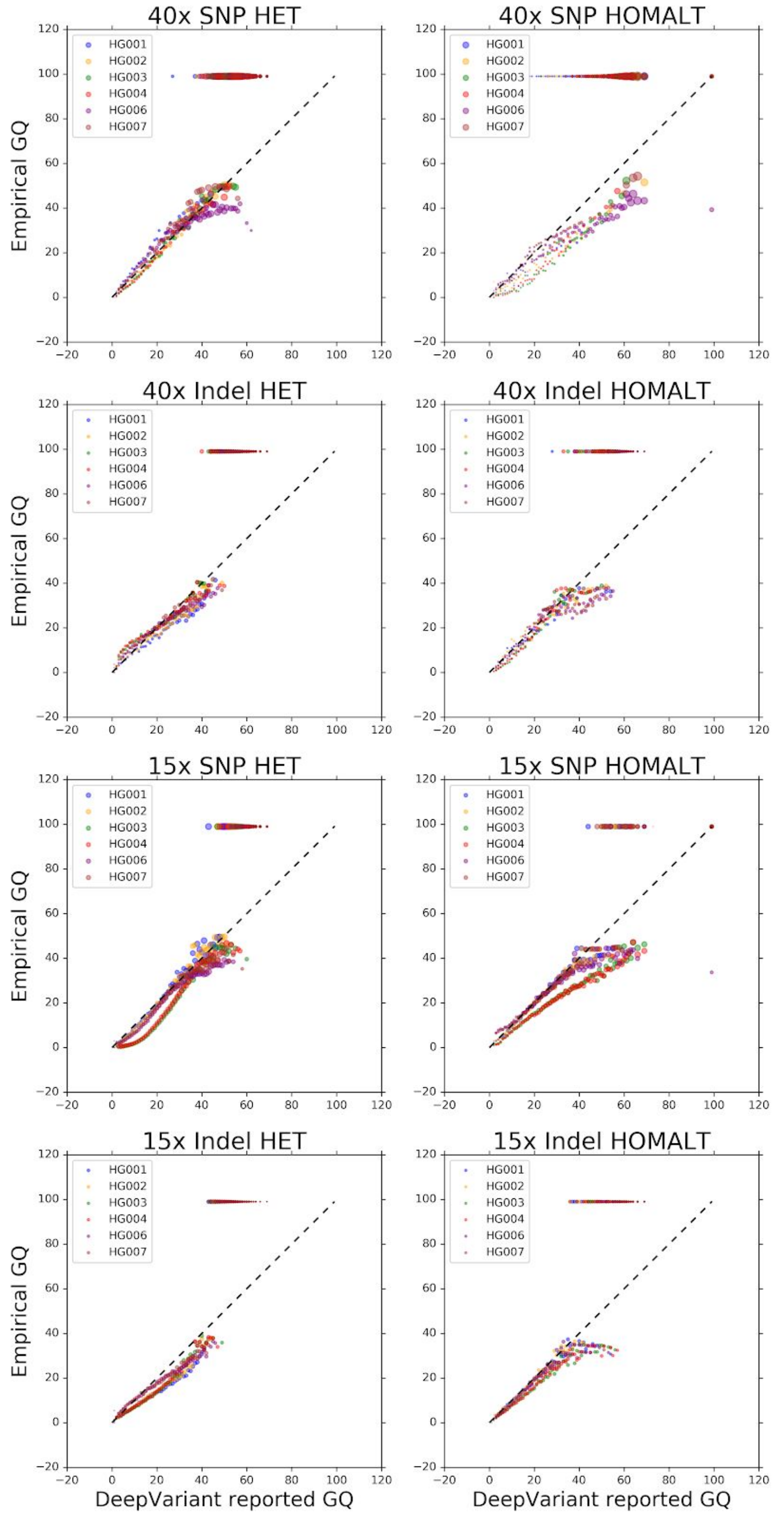


**Supplementary Figure 2. DeepVariant 0.8 genotype quality (GQ) score calibration stratified by variant type.** Similar to Figure 1, for both ~40x and 15x coverage reads and computed separately per variant type (SNP, indel) and zygosity (heterozygous reference/alternate, homozygous alternate).

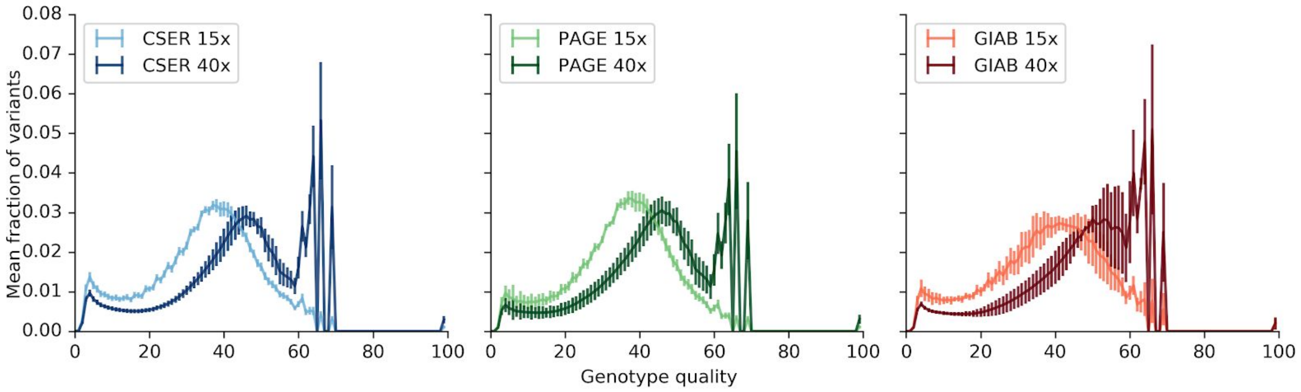


**Supplementary Figure 3. Sample genotype quality distributions for DeepVariant v0.8.0 calls as a function of sequence coverage.** For each of the three development datasets, average fractions of variants at each estimated genotype quality are plotted at both 15x and 40x sequence coverage. Error bars represent sample standard deviations.

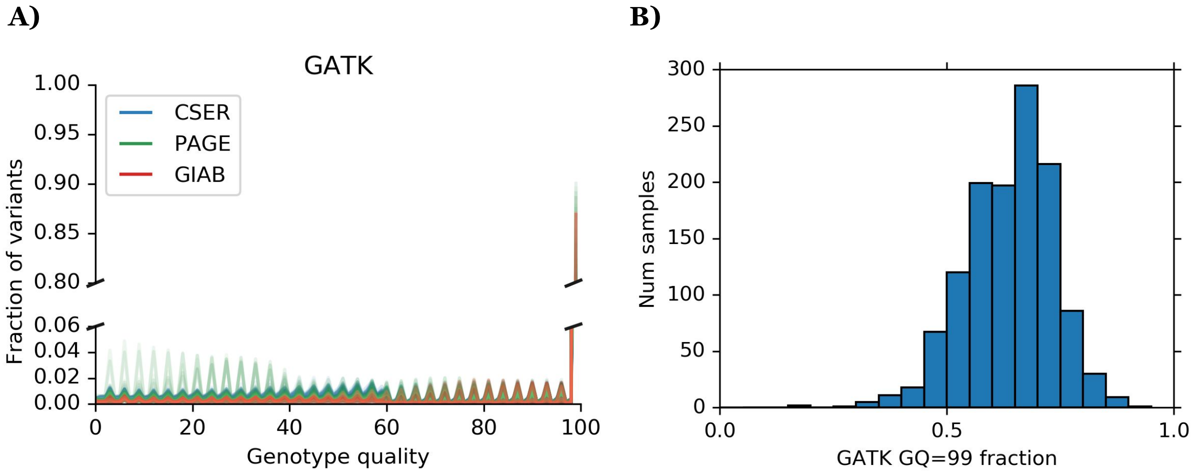


**Supplementary Figure 4. Genotype quality (GQ) distribution properties of GATK PASS variants. A)** Distribution of reported GQ for GATK HaplotypeCaller in all 1,248 samples from GIAB, PAGE, and CSER computed on chromosome 2. Note the broken y-axis and different scales. **B)** The fraction of variant calls with GQ=99 from GATK4 HaplotypeCaller across the 1,248 samples. On average, 63.81% of variants have GQ=99.

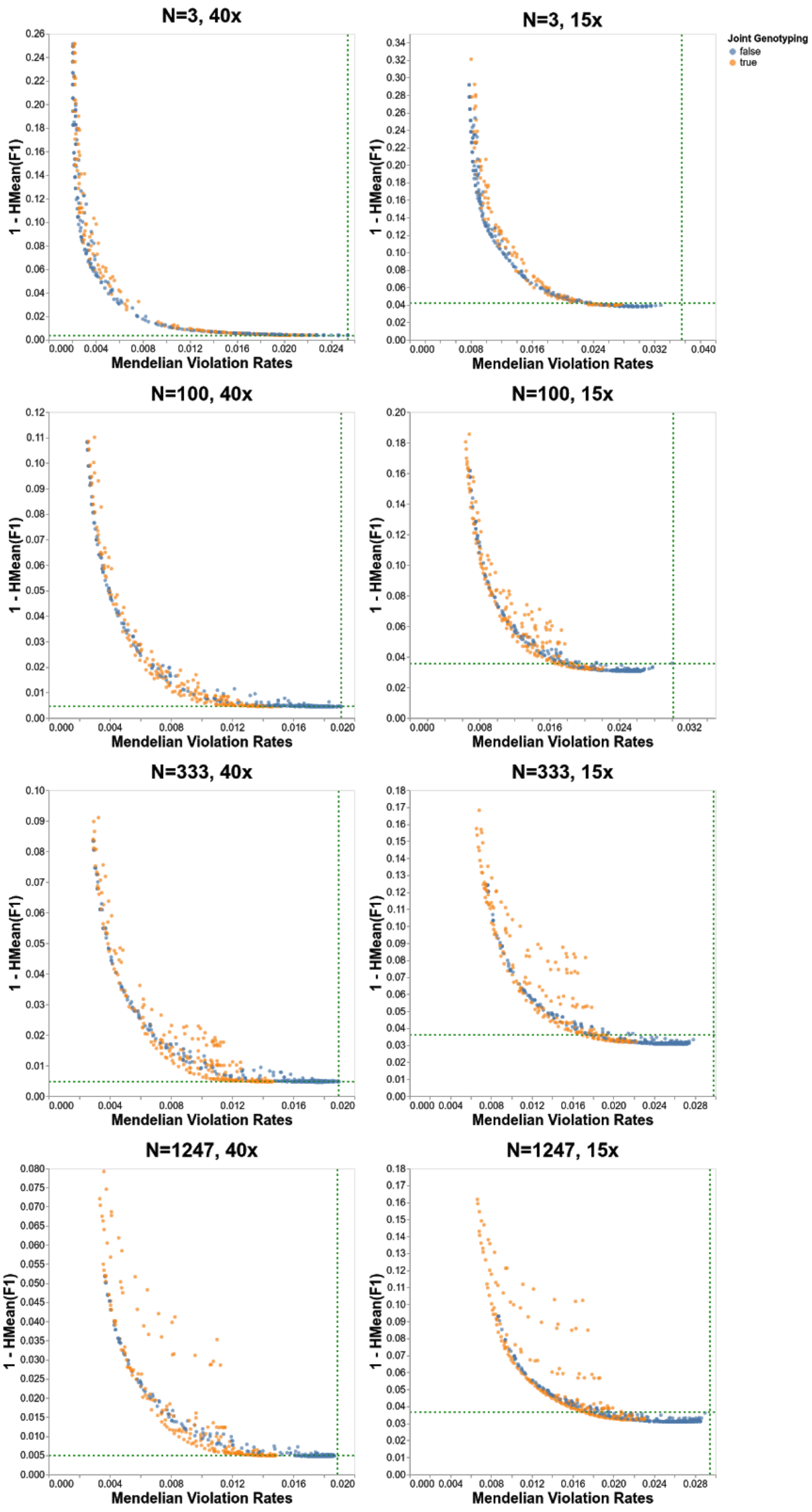


**Supplementary Figure 5. Pareto-optimal search for all WGS cohorts.** See also **Figure 2**.

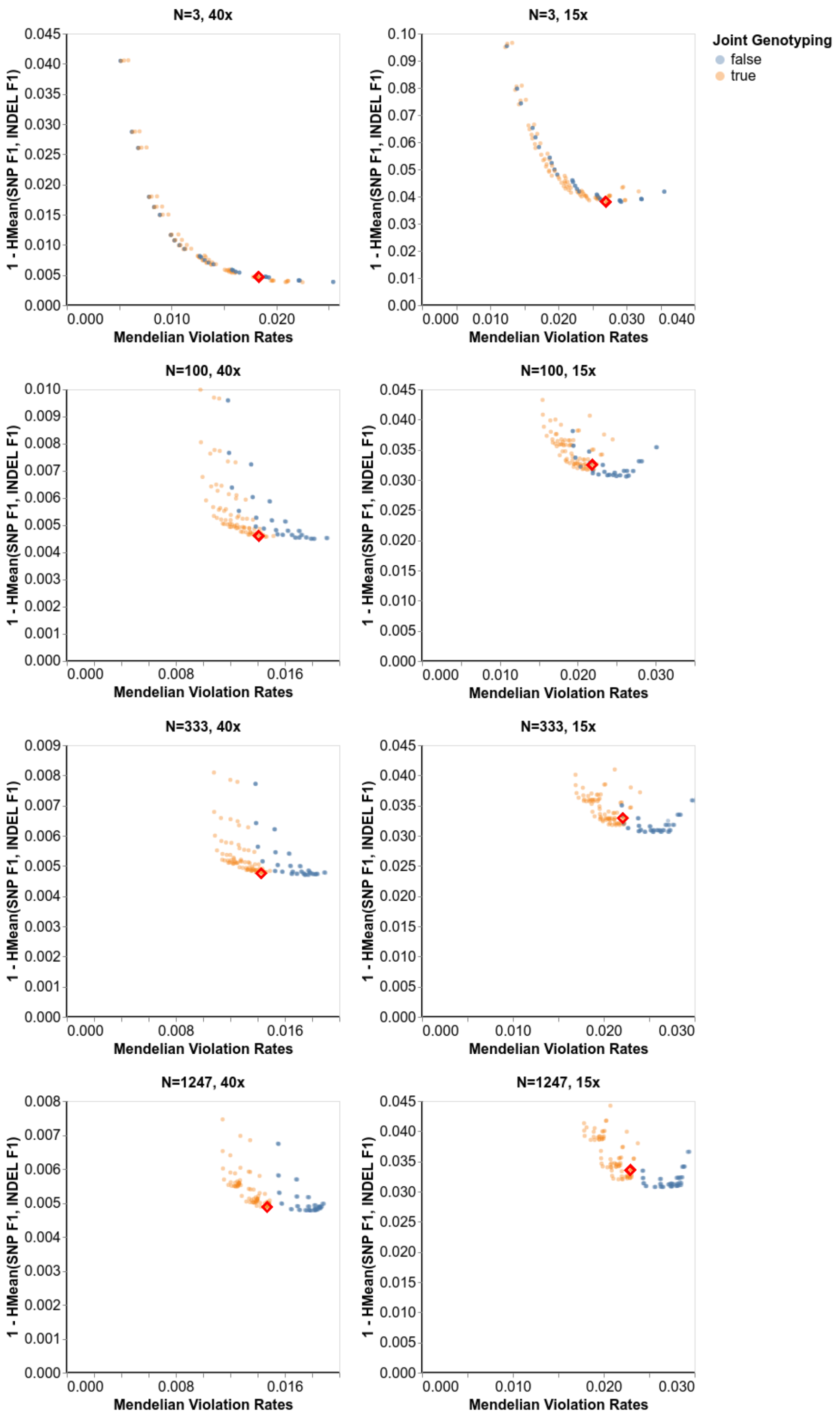


**Supplementary Figure 6. Grid search for GLnexus parameters.** Each data point represents a unique parameter combination. The x-axis shows the rates of Mendelian violations and the y-axis shows one minus the harmonic mean of SNP F1 and indel F1 using GIAB benchmark calls (lower numbers are better). The red highlighted points are the optimized parameter set used for DV-GLN-OPT. See also **Supplementary Figures 7** and **8**.

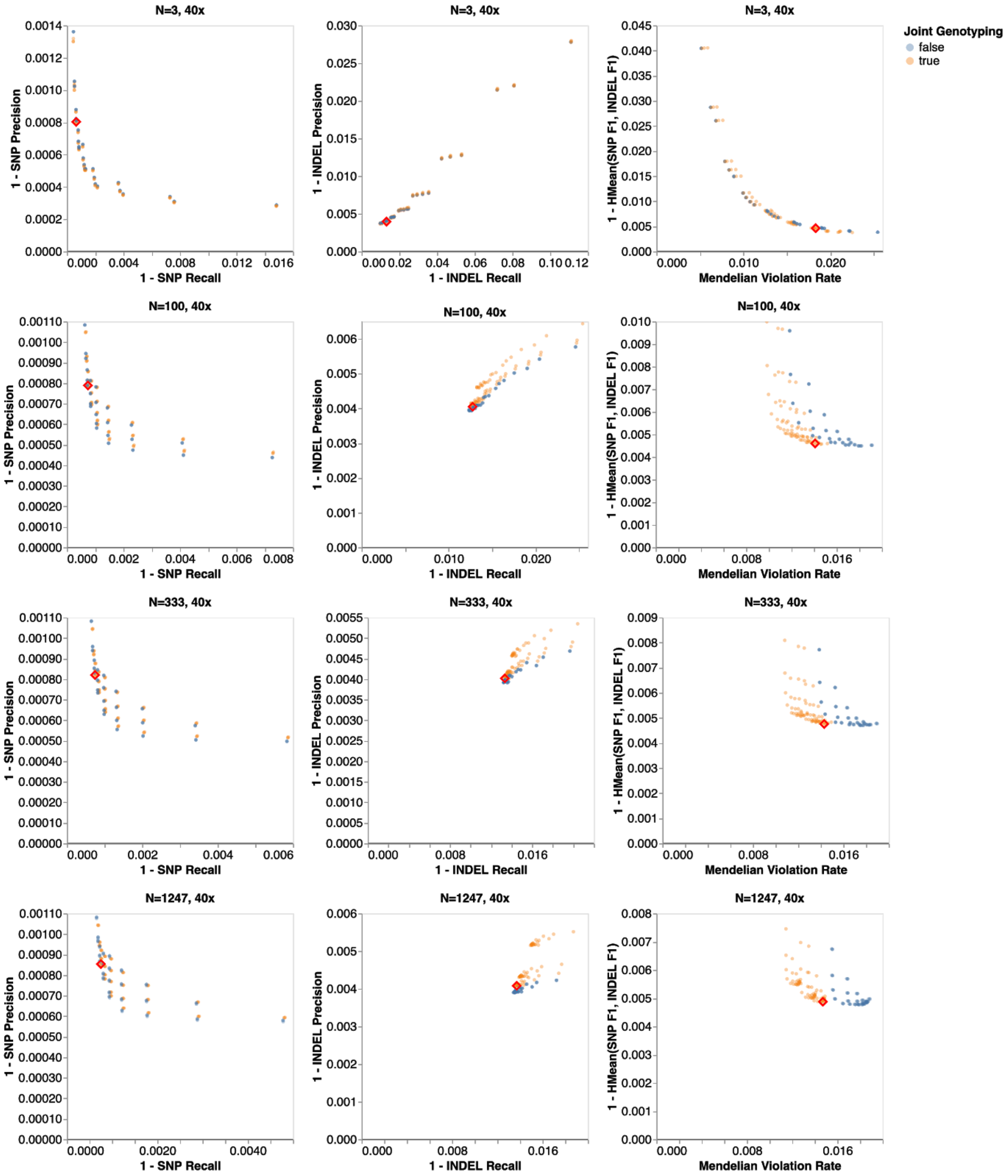


**Supplementary Figure 7. Optimized parameter performance for all 40x sequence coverage cohorts, compared to other parameter sets explored by grid search.** The red highlighted points are the optimized parameter set.

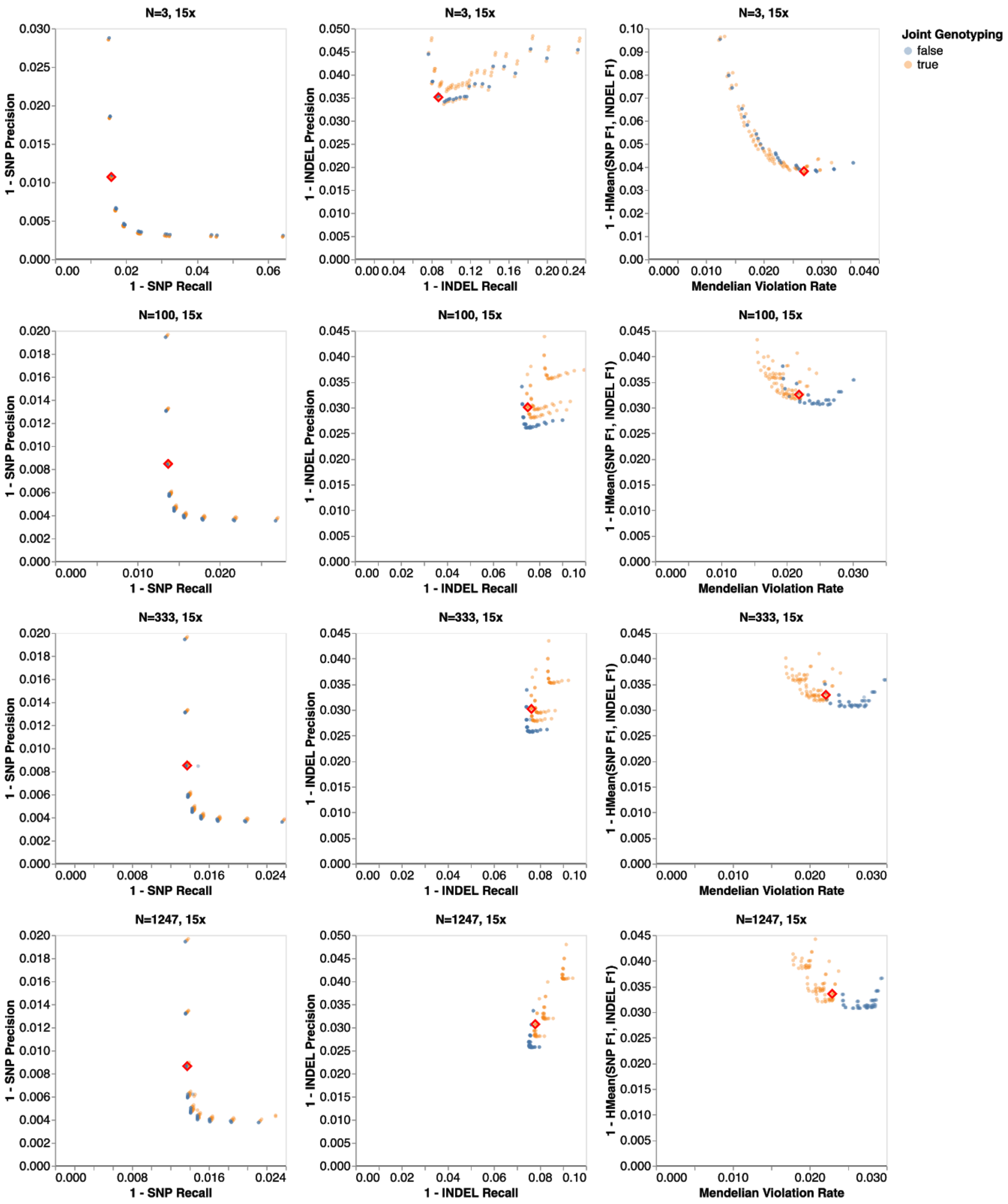


**Supplementary Figure 8. Optimized parameter performance for all 15x sequence coverage cohorts, compared to other parameter sets explored by grid search.** The red highlighted points are the optimized parameter set.

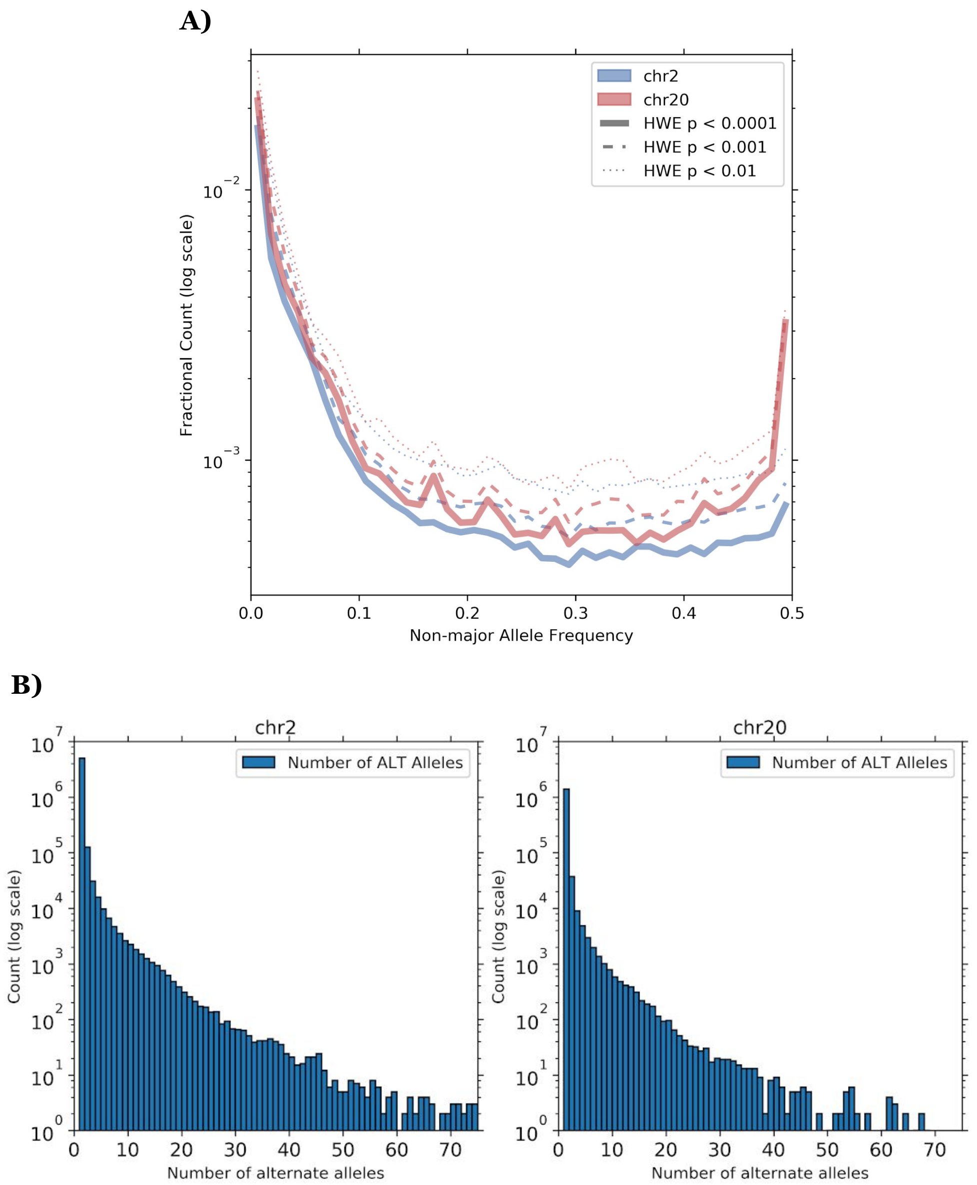


**Supplementary Figure 9. Similarity of calls in chr2 and chr20 of N=1,247 cohort with 40x coverage generated by the optimized DeepVariant+GLnexus pipeline. A)** Fractional counts of variants with low HWE p-values, binned by non-major allele frequency in chromosome 2 and chromosome 20. **B)** Histogram of the number of alternate alleles in variants in chr2 and chr20. Note: chr2 is ~4x larger than chr20.

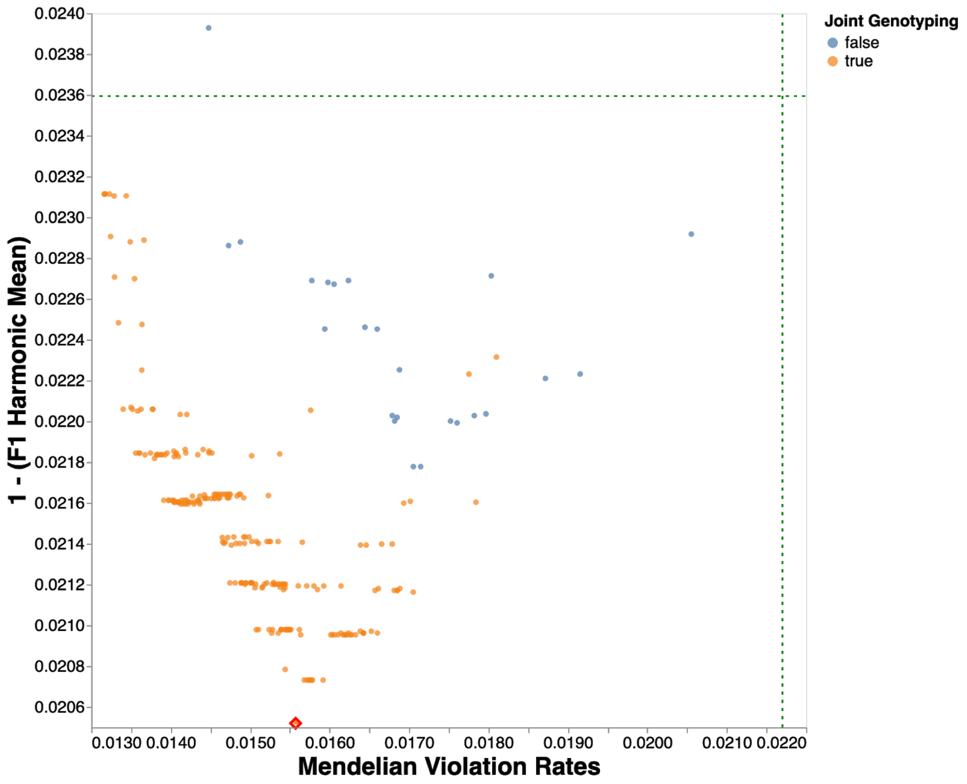


**Supplementary Figure 10. Pareto optimal search for exomes.** The red diamond indicates the exome-optimized DeepVariant+GLnexus pipeline.

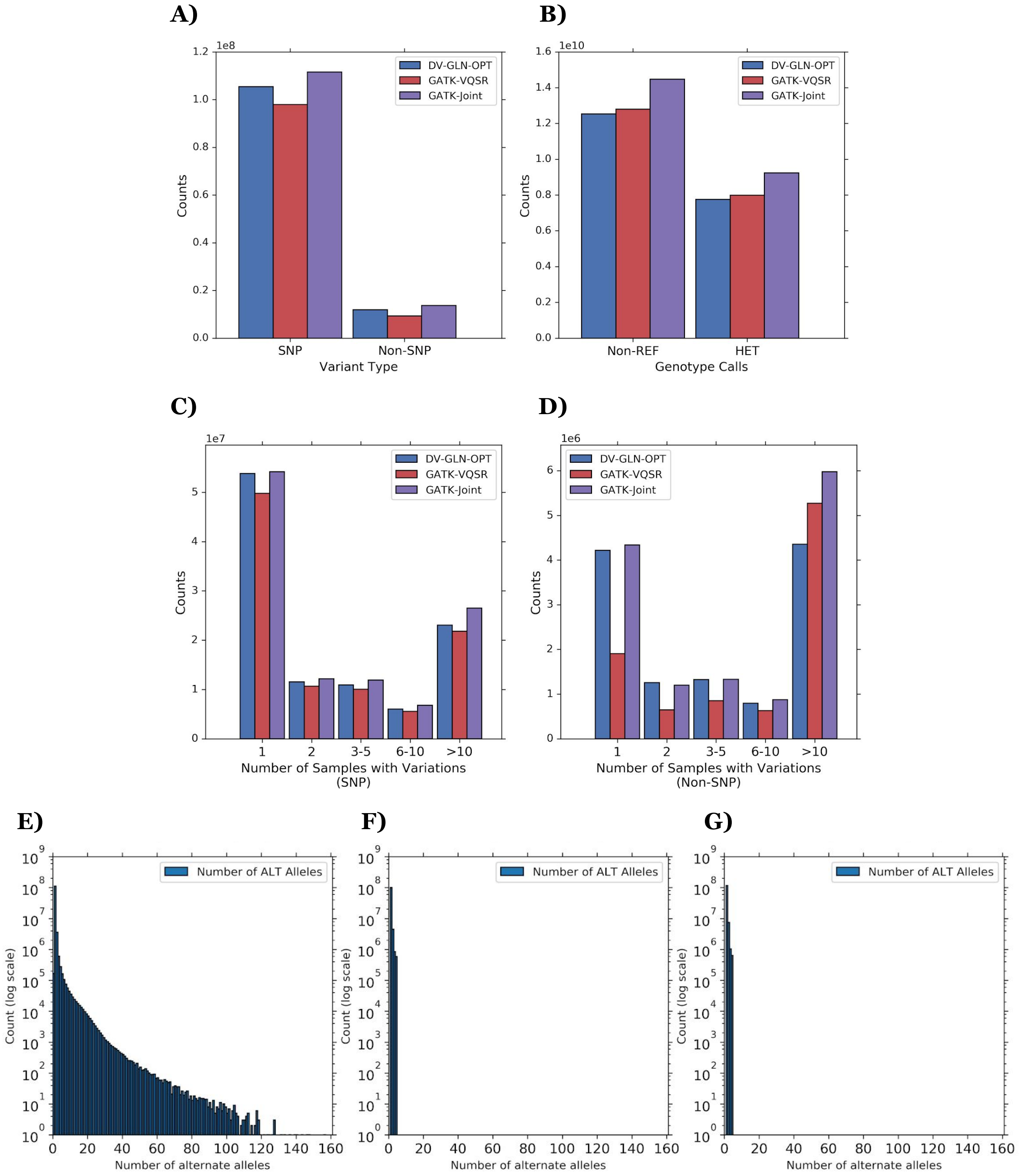


**Supplementary Figure 11**. **Comparison of 1KGP cohort callset properties.**
**A)** The number of variants generated by each method per variant type. **B)** The number of all genotype calls excluding homozygous reference calls (Non-REF), and the number of heterozygous genotype calls in each method. **C, D)** Histogram of number of samples with variations in SNPs (C) and non-SNPs (D). Per each cohort variant (i.e. a row in a cohort VCF) of each type, we count the number of samples with a non-reference genotype (i.e. the leftmost bins are singletons). **E, F, G)** Histogram of the count of the number of alternate alleles from DV-GLN-OPT (E), GATK-VQSR (F), and GATK-Joint (G). Note that GATK limits the number of alternate alleles to 6 by default (gatk.broadinstitute.org/hc/en-us/articles/360036734631-GenotypeGVCFs).

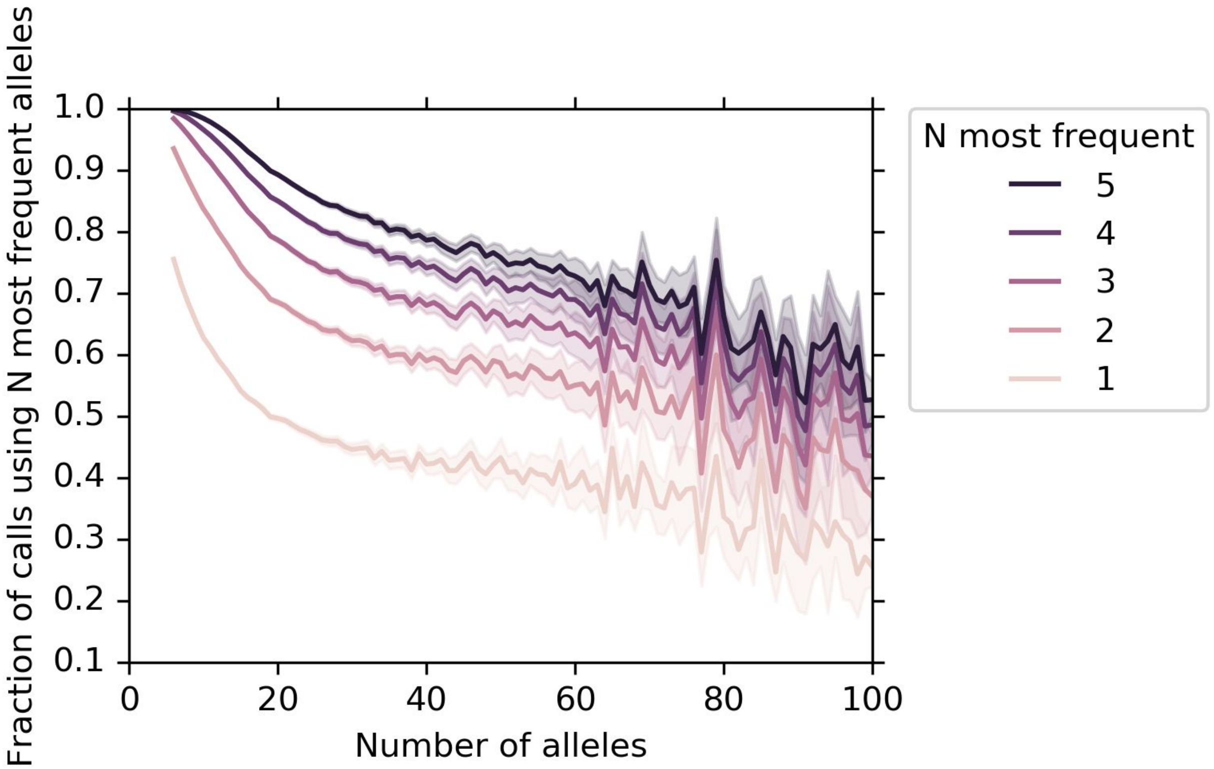


**Supplementary Figure 12. Distribution of allele usage in DeepVariant multiallelic sites.** All variants in the 1KGP cohort with at least six total alleles were analyzed to see the fraction of calls using the most frequently-called alleles. The "100" allele bin contains all variants (n=97) with 100 or more alleles (the maximal variant contained 162 alleles). Bands represent 95% confidence intervals.

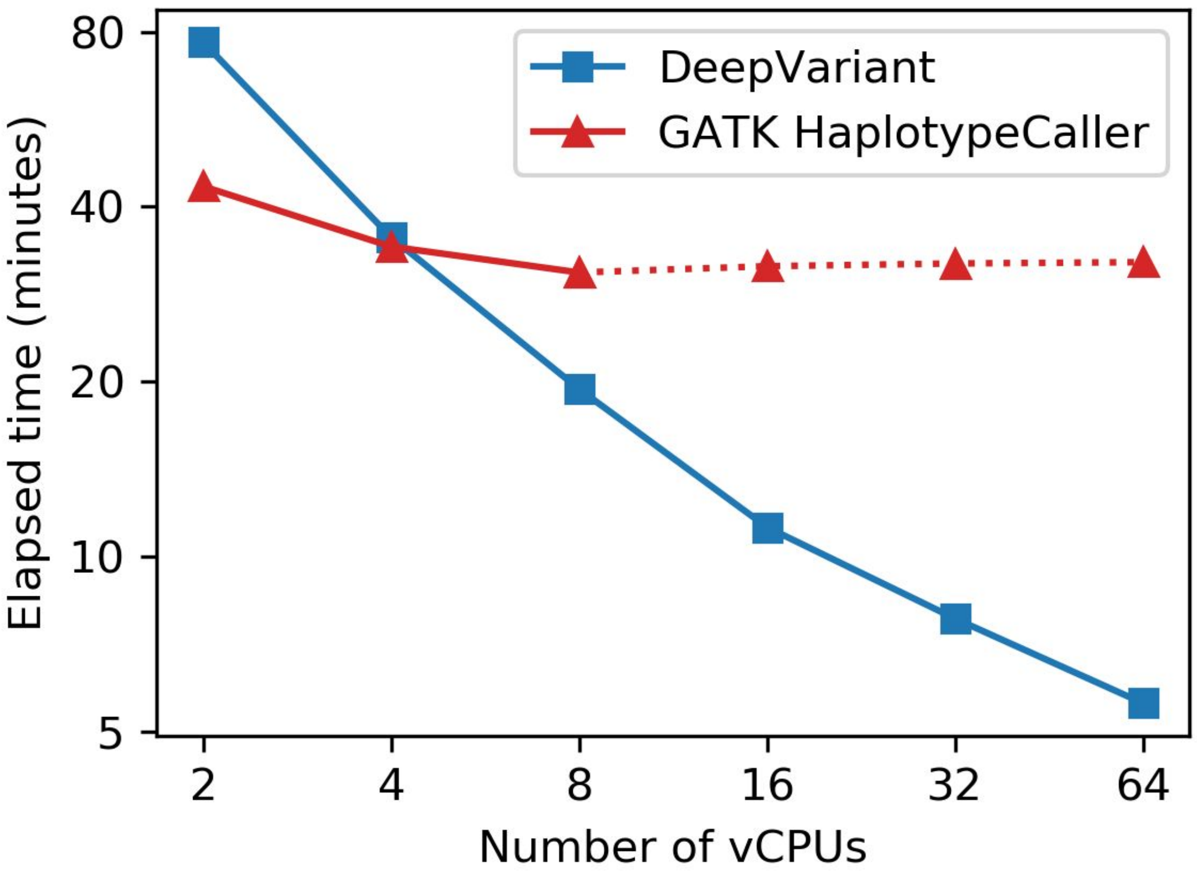


**Supplementary Figure 13. Log-scaled elapsed real times to generate chr22 gVCF of one sample (NA12878) using a varying number of vCPUs.** Identical to **Figure 7B**, but using log scales for both axes.

### Supplementary Tables

**Supplementary Table 1: Cohort evaluation experimental setup.**

**A)** Cohort subset definitions.

| Source | Subset Name | Size | Description |
|---|---|---|---|
| GIAB | *GIAB3* | 3 | HG002, HG003, HG004 (son, father, mother) trio. |
| | *GIAB5* | 5 | HG001, HG003, HG004, HG006, HG007 (mutually non-descendant samples). |
| | *GIAB_WES* | 2 | HG001, HG002 exomes sequences. |
| CSER | *CSER15* | 15 | Randomly selected 5 WGS trios, excluding outliers. See **Supplementary Table 2**. |
| | *CSER* | 929 | All available WGS CSER samples. |
| | *CSER15_WES* | 15 | Randomly selected 5 WES trios, excluding outliers. See **Supplementary Table 2**. |
| | *CSER_WES* | 344 | All available WES CSER samples. |
| PAGE | *PAGE80* | 80 | Randomly selected 80 PAGE samples, excluding outliers. See **Supplementary Table 5**. |
| | *PAGE* | 313 | All PAGE samples. |

**B)** Custom cohorts for cohort evaluation. Each WGS cohort has two versions for 40x and 15x coverage.

| Cohort Name | Size | Definition | Single-sample benchmark samples | Evaluation trios (for Mendelian violation) |
|---|---|---|---|---|
| *GIAB3* | 3 | GIAB3 | GIAB3 | GIAB3 |
| *GIAB5_CSER15_PAGE80* | 100 | GIAB5 + CSER15 + PAGE80 | GIAB5 | CSER15 |
| *GIAB5_CSER15_PAGE* | 333 | GIAB5 + CSER15 + PAGE | GIAB5 | CSER15 |
| *GIAB5_CSER_PAGE* | 1,247 | GIAB5 + CSER + PAGE | GIAB5 | CSER15 |
| *GIAB_CSER_WES* | 346 | GIAB_WES + CSER_WES | GIAB_WES | CSER15_WES |

**C)** Cohort evaluation metrics.

| **Cohort Metric Name** | **Definition** |
|---|---|
| Mendelian Violation | Arithmetic mean of Mendelian violation rates on evaluation trios. |
| SNP Precision | Arithmetic mean of SNP precisions of all single-sample benchmark samples. |
| SNP Recall | Arithmetic mean of SNP recalls of all single-sample benchmark samples. |
| Indel Precision | Arithmetic mean of indel precisions of all single-sample benchmark samples. |
| Indel Recall | Arithmetic mean of indel recalls of all single-sample benchmark samples. |

**Supplementary Table 2: CSER15 and CSER15_WES trio sample names.**

| | **Child** | **Father** | **Mother** |
|---|---|---|---|
| **CSER15** | SRR4370493 | SRR4370494 | SRR4370495 |
| | SRR6706862 | SRR6707105 | SRR6707106 |
| | SRR6706955 | SRR6706956 | SRR6706957 |
| | SRR6707156 | SRR6707157 | SRR6707158 |
| | SRR6707268 | SRR6707269 | SRR6707270 |
| **CSER15_WES** | SRR3406206 | SRR3406207 | SRR3406279 |
| | SRR3406280 | SRR3406209 | SRR3406430 |
| | SRR3406315 | SRR3406316 | SRR3406317 |
| | SRR3406410 | SRR3406404 | SRR3406373 |
| | SRR3406427 | SRR3406428 | SRR3406429 |

**Supplementary Table 3: GLnexus configurable parameters.**

| Name | Type | Default Value | Tuned | Description |
|---|---|---|---|---|
| `min_AQ1` | Numeric | 0 | Y | The minimum allele quality in phred scale to be used for all alleles. Alleles lower than this quality will be pruned. Increasing this will increase specificity and decrease sensitivity. |
| `min_AQ2` | Numeric | 0 | Y | The minimum allele quality in phred scale to be used for alleles that have multiple observations. min_AQ1 ≥ min_AQ2. |
| `min_GQ` | Numeric | 0 | Y | The minimum genotype quality in phred scale to be used for copy number estimates for the constituent alleles. |
| `min_allele_copy_number` | Numeric | 1 | | The minimum number of observations an allele needs to have in order to be kept. |
| `revise_genotypes` | Boolean | false | Y | If true, joint calling is enabled - use genotype likelihoods and allele frequencies to revise low quality genotype calls. |
| `min_assumed_allele_frequency` | Numeric | 0.0001 | | Allele frequency lower than this value will be fixed to be this minimum value so rare alleles are less likely to be lost in a large cohort. |
| `required_dp` | Numeric | 0 | | The minimum depth required for any allele call. |

**Supplementary Table 4. Comparison of variant calling-merging methods.**
**A)** GATK-Joint, GATK-VQSR, DV-GLN-NOMOD, and DV-GLN-OPT pipelines were compared by GIAB sample concordance and trio sample Mendelian violation rates for all 40x sequence coverage cohorts. All evaluation metrics were computed on chromosome 20. Bold numbers are the best values across three methods, or the values that are within 0.001 difference from the best value. The parameters and resources used for GATK-VQSR can be found in **Supplementary Note 3**. GATK-VQSR is skipped for the trio cohort due to the insufficient size. The trio cohort also includes a cohort generated from DeepVariant single-sample calls merged using GATK GenotypeGVCFs ("DV-GATK"). Prec, precision; MV, Mendelian violation rate; Std, standard deviation. **B)** Similar to A), for the 15x sequence coverage cohorts. **C)** Similar to A) and B), for a single 346-individual exome cohort. A separate parameter set "OPT-WES" for DV+GLnexus, optimized specifically for exomes, is used.

**A)** 40x coverage

| Size | Method | SNP F1 | SNP Recall | SNP Recall Std | SNP Prec | SNP Prec Std | Indel F1 | Indel Recall | Indel Recall Std | Indel Prec | Indel Prec Std | MVR | MVR Std |
|---|---|---|---|---|---|---|---|---|---|---|---|---|---|
| **3** | GATK (Joint) | 0.99788 | **0.99945** | 0.00013 | 0.99631 | 0.00033 | 0.97950 | 0.97285 | 0.01233 | 0.98624 | 0.00551 | 6.61% | . |
| | DV-GATK (Joint) | 0.97547 | 0.98700 | 0.00787 | 0.96420 | 0.02269 | 0.92497 | 0.90430 | 0.04933 | 0.94660 | 0.03019 | **1.66%** | . |
| | DV-GLN (NOMOD) | **0.99936** | **0.99937** | 0.00019 | **0.99936** | 0.00025 | **0.99036** | **0.98556** | 0.00685 | **0.99521** | 0.00131 | 4.72% | . |
| | DV-GLN (OPT) | **0.99932** | **0.99903** | 0.00011 | **0.99962** | 0.00013 | 0.98753 | 0.98057 | 0.00988 | **0.99459** | 0.00193 | 3.32% | . |
| **100** | GATK (Joint) | 0.99657 | **0.99916** | 0.00057 | 0.99399 | 0.00205 | 0.97858 | 0.97727 | 0.01019 | 0.97989 | 0.00502 | 5.97% | 0.26% |
| | GATK (VQSR) | 0.98638 | 0.97429 | 0.00145 | 0.99877 | 0.00071 | 0.97175 | 0.96146 | 0.01064 | 0.98226 | 0.00614 | 4.15% | 0.30% |
| | DV-GLN (NOMOD) | **0.99935** | **0.99929** | 0.00021 | **0.99941** | 0.00024 | **0.98959** | **0.98429** | 0.00727 | **0.99495** | 0.00065 | 2.46% | 0.21% |
| | DV-GLN (OPT) | **0.99934** | **0.99917** | 0.00023 | **0.99951** | 0.00015 | **0.98926** | **0.98387** | 0.00698 | **0.99472** | 0.00031 | **1.61%** | 0.19% |
| **333** | GATK (Joint) | 0.99634 | **0.99915** | 0.00056 | 0.99355 | 0.00234 | 0.97745 | 0.97633 | 0.01060 | 0.97858 | 0.00487 | 6.37% | 0.27% |
| | GATK (VQSR) | 0.98886 | 0.97935 | 0.00135 | 0.99855 | 0.00074 | 0.97156 | 0.96195 | 0.01126 | 0.98137 | 0.00625 | 4.42% | 0.30% |
| | DV-GLN (NOMOD) | **0.99932** | **0.99922** | 0.00024 | **0.99942** | 0.00025 | **0.98907** | **0.98334** | 0.00790 | **0.99487** | 0.00069 | 2.44% | 0.20% |
| | DV-GLN (OPT) | **0.99931** | **0.99910** | 0.00027 | **0.99951** | 0.00017 | **0.98903** | **0.98332** | 0.00741 | **0.99480** | 0.00039 | **1.63%** | 0.19% |
| **1247** | GATK (Joint) | 0.99609 | **0.99912** | 0.00057 | 0.99309 | 0.00264 | 0.97666 | 0.97523 | 0.01057 | 0.97811 | 0.00540 | 7.03% | 0.29% |
| | GATK (VQSR) | 0.98771 | 0.97709 | 0.00526 | **0.99857** | 0.00072 | 0.97081 | 0.96101 | 0.01148 | 0.98082 | 0.00636 | 5.00% | 0.33% |
| | DV-GLN (NOMOD) | **0.99931** | **0.99922** | 0.00025 | **0.99941** | 0.00025 | **0.98868** | **0.98245** | 0.00855 | **0.99500** | 0.00065 | 2.43% | 0.20% |
| | DV-GLN (OPT) | **0.99930** | **0.99913** | 0.00028 | **0.99948** | 0.00017 | **0.98860** | **0.98261** | 0.00806 | **0.99466** | 0.00047 | **1.69%** | 0.20% |

**B)** 15x coverage

| Size | Method | SNP F1 | SNP Recall | SNP Recall Std | SNP Prec | SNP Prec Std | Indel F1 | Indel Recall | Indel Recall Std | Indel Prec | Indel Prec Std | MVR | MVR Std |
|---|---|---|---|---|---|---|---|---|---|---|---|---|---|
| **3** | GATK (Joint) | 0.98176 | 0.97426 | 0.00274 | **0.98937** | 0.00284 | 0.88575 | 0.84140 | 0.03604 | 0.93503 | 0.01991 | 8.64% | . |
| | DV-GATK (Joint) | 0.96691 | 0.94780 | 0.02786 | 0.98680 | 0.00564 | 0.86053 | 0.80400 | 0.05586 | 0.92560 | 0.02171 | **2.39%** | . |
| | DV-GLN (NOMOD) | 0.97596 | **0.98108** | 0.00198 | 0.97090 | 0.01670 | **0.92567** | **0.90176** | 0.01695 | 0.95090 | 0.01376 | 5.20% | . |
| | DV-GLN (OPT) | **0.98448** | **0.98043** | 0.00230 | **0.98857** | 0.00590 | **0.92479** | 0.89072 | 0.02129 | **0.96158** | 0.00667 | 3.81% | . |
| **100** | GATK (Joint) | 0.98498 | 0.98079 | 0.00584 | 0.98920 | 0.00205 | 0.91734 | 0.88600 | 0.04947 | 0.95099 | 0.01625 | 8.25% | 0.31% |
| | GATK (VQSR) | 0.97231 | 0.95344 | 0.00759 | **0.99194** | 0.00179 | 0.91038 | 0.87252 | 0.04989 | 0.95168 | 0.01729 | 6.93% | 0.50% |
| | DV-GLN (NOMOD) | 0.98260 | **0.98479** | 0.00461 | 0.98042 | 0.01743 | 0.93578 | **0.91087** | 0.02878 | 0.96210 | 0.01692 | 3.46% | 0.25% |
| | DV-GLN (OPT) | **0.98789** | **0.98453** | 0.00417 | **0.99128** | 0.00562 | **0.93720** | 0.90884 | 0.02721 | **0.96738** | 0.00972 | **2.33%** | 0.16% |
| **333** | GATK (Joint) | 0.98480 | 0.98079 | 0.00578 | 0.98885 | 0.00245 | 0.91739 | 0.88560 | 0.04938 | 0.95156 | 0.01514 | 8.66% | 0.29% |
| | GATK (VQSR) | 0.97298 | 0.95401 | 0.00722 | **0.99271** | 0.00183 | 0.91152 | 0.87378 | 0.05004 | 0.95266 | 0.01643 | 7.18% | 0.47% |
| | DV-GLN (NOMOD) | 0.98254 | **0.98466** | 0.00467 | 0.98043 | 0.01743 | 0.93494 | **0.90900** | 0.02968 | 0.96241 | 0.01679 | 3.43% | 0.25% |
| | DV-GLN (OPT) | **0.98780** | **0.98444** | 0.00419 | 0.99120 | 0.00565 | **0.93655** | 0.90755 | 0.02804 | **0.96746** | 0.00979 | **2.37%** | 0.15% |
| **1247** | GATK (Joint) | 0.98447 | 0.98082 | 0.00575 | 0.98815 | 0.00284 | 0.91630 | 0.88610 | 0.04937 | 0.94863 | 0.01606 | 9.37% | 0.25% |
| | GATK (VQSR) | 0.97513 | 0.95795 | 0.00971 | **0.99293** | 0.00155 | 0.91099 | 0.87506 | 0.04970 | 0.95001 | 0.01698 | 8.06% | 0.47% |
| | DV-GLN (NOMOD) | 0.98248 | **0.98454** | 0.00474 | 0.98043 | 0.01743 | 0.93324 | **0.90532** | 0.03144 | 0.96294 | 0.01651 | 3.38% | 0.25% |
| | DV-GLN (OPT) | **0.98773** | **0.98440** | 0.00416 | 0.99108 | 0.00572 | **0.93506** | **0.90492** | 0.02936 | **0.96727** | 0.00945 | **2.46%** | 0.16% |

**C)** Exome

| Size | Method | SNP F1 | SNP Recall | SNP Recall Std | SNP Prec | SNP Prec Std | INDEL F1 | INDEL Recall | INDEL Recall Std | INDEL Prec | INDEL Prec Std | MVR | MVR Std |
|---|---|---|---|---|---|---|---|---|---|---|---|---|---|
| **346** | GATK (Joint) | 0.98918 | **0.99409** | 0.00482 | 0.98433 | 0.00940 | 0.81343 | 0.91792 | 0.03306 | 0.73030 | 0.11674 | 2.69% | 0.29% |
| | GATK (VQSR) | 0.99122 | 0.99318 | 0.00472 | 0.98928 | 0.00825 | 0.81945 | 0.90645 | 0.03309 | 0.74769 | 0.11412 | 2.16% | 0.21% |
| | DV-GLN (NOMOD) | **0.99591** | **0.99464** | 0.00383 | **0.99718** | 0.00058 | 0.95813 | 0.93523 | 0.00595 | 0.98218 | 0.00571 | 2.15% | 0.36% |
| | DV-GLN (OPT-WES) | **0.99600** | **0.99468** | 0.00380 | **0.99732** | 0.00050 | **0.96356** | **0.94096** | 0.00839 | **0.98727** | 0.00408 | **1.56%** | 0.20% |

**Supplementary Table 5. Imputation accuracy of GIAB benchmark callsets.** The imputed variant calls of HG002 and HG005 are scored using the GIAB benchmark variants v3.3.2 (GRCh38) and hap.py v0.3.9. Two evaluation regions are used: "*full conf. region*" is the intersection of the HG002 and HG005 benchmark regions, agnostic to either reference panel, and "*shared conf. region in both panels*" is the subset of full conf. region that also intersects both the DV-GLN-OPT panel and GATK panel regions.

| Eval. region | Sample | Ref. panel method | Type | F1 | Recall | Precision | TP | FN | FP | FP.gt |
|---|---|---|---|---|---|---|---|---|---|---|
| Full conf. region | HG002 | DV-GLN-OPT | INDEL | **0.90307** | **0.88392** | **0.92308** | 325366 | 42729 | 27115 | 14189 |
| | | | SNP | **0.94555** | **0.92818** | **0.96359** | 2596802 | 200944 | 98143 | 39149 |
| | | GATK | INDEL | 0.89921 | 0.87839 | 0.92106 | 323329 | 44766 | 27721 | 14140 |
| | | | SNP | 0.94219 | 0.92176 | 0.96354 | 2578852 | 218894 | 97606 | 38968 |
| | HG005 | DV-GLN-OPT | INDEL | **0.89325** | **0.88232** | **0.90446** | 319121 | 42564 | 33714 | 15193 |
| | | | SNP | **0.93832** | **0.93058** | 0.94618 | 2566405 | 191442 | 146023 | 41306 |
| | | GATK | INDEL | 0.88989 | 0.87706 | 0.90310 | 317221 | 44468 | 34052 | 15051 |
| | | | SNP | 0.93511 | 0.92425 | **0.94624** | 2548940 | 208921 | 144832 | 41045 |
| Shared conf. region in both panels | HG002 | DV-GLN-OPT | INDEL | **0.90621** | 0.88959 | **0.92345** | 323048 | 40094 | 26778 | 14018 |
| | | | SNP | **0.95033** | **0.93721** | **0.96382** | 2579381 | 172822 | 96838 | 38671 |
| | | GATK | INDEL | 0.90549 | **0.88971** | 0.92183 | 323092 | 40050 | 27399 | 14126 |
| | | | SNP | 0.95016 | 0.93696 | 0.96373 | 2578713 | 173490 | 97058 | 38965 |
| | HG005 | DV-GLN-OPT | INDEL | **0.89633** | 0.88827 | **0.90455** | 316799 | 39850 | 33430 | 15031 |
| | | | SNP | **0.94287** | **0.93947** | 0.94630 | 2549478 | 164268 | 144708 | 40856 |
| | | GATK | INDEL | 0.89593 | **0.88863** | 0.90336 | 316929 | 39720 | 33907 | 15032 |
| | | | SNP | 0.94276 | 0.93917 | **0.94638** | 2548675 | 165071 | 144442 | 41046 |

**Supplementary Table 6. DeepVariant and GATK HaplotypeCaller benchmark.** Summary statistics of elapsed real time, user CPU time, and system CPU time spent on running DeepVariant and GATK HaplotypeCaller across 2,504 1KGP samples, chromosome 22 only, using 8-vCPU virtual machines.

| | **DeepVariant** (seconds) | | | **GATK HaplotypeCaller** (seconds) | | |
|---|---|---|---|---|---|---|
| | **Real** | **User CPU** | **System CPU** | **Real** | **User CPU** | **System CPU** |
| **Mean** | 1,201.26 | 7,733.31 | 146.56 | 1,989.20 | 5,346.24 | 9.63 |
| **St. dev.** | 75.44 | 470.21 | 10.20 | 203.66 | 511.87 | 1.48 |
| **Min** | 1,044.64 | 6,795.50 | 125.96 | 1,617.60 | 4,442.35 | 7.59 |
| **25%** | 1,144.85 | 7,386.25 | 138.91 | 1,849.41 | 4,986.84 | 8.51 |
| **50%** | 1,178.51 | 7,594.58 | 143.52 | 1,953.21 | 5,223.30 | 9.09 |
| **75%** | 1,263.78 | 8,113.92 | 154.78 | 2,082.78 | 5,636.03 | 10.66 |
| **Max** | 1,484.26 | 9,279.03 | 191.90 | 3,601.13 | 9,196.31 | 18.43 |

**Supplementary Table 7. PAGE80 sample names.**

| | | | | |
|---|---|---|---|---|
| SRR2993850 | SRR2994215 | SRR2994285 | SRR2994293 | SRR2994301 |
| SRR2994861 | SRR2995075 | SRR2995970 | SRR2996055 | SRR2996085 |
| SRR2996123 | SRR2996131 | SRR2996243 | SRR2996321 | SRR2996337 |
| SRR2996373 | SRR3003654 | SRR3003716 | SRR3003902 | SRR3004018 |
| SRR3004154 | SRR3004266 | SRR3010823 | SRR3010896 | SRR3010944 |
| SRR3011061 | SRR3011110 | SRR3011469 | SRR3011551 | SRR3011961 |
| SRR3012267 | SRR3012323 | SRR3012447 | SRR3012511 | SRR3012726 |
| SRR3012734 | SRR3012758 | SRR3012834 | SRR3012951 | SRR3012975 |
| SRR3013049 | SRR3013065 | SRR3013089 | SRR3013153 | SRR3013161 |
| SRR3013177 | SRR3013201 | SRR3013202 | SRR3013242 | SRR3013338 |
| SRR3013370 | SRR3013378 | SRR3013430 | SRR3013508 | SRR3013524 |
| SRR3013587 | SRR3013603 | SRR3013793 | SRR3013843 | SRR3013881 |
| SRR3014027 | SRR3014035 | SRR3014051 | SRR3014088 | SRR3014096 |
| SRR3014120 | SRR3014152 | SRR3014168 | SRR3014200 | SRR3014306 |
| SRR3014314 | SRR3014338 | SRR3014370 | SRR3014378 | SRR3014418 |
| SRR3014442 | SRR3014520 | SRR3014536 | SRR3014653 | SRR3014824 |

### Supplementary Notes

#### Supplementary Note 1: "DV-GLN-OPT" optimized GLnexus WGS configuration

This configuration is available as “DeepVariantWGS” in GLnexus v1.2.2: https://github.com/dnanexus-rnd/GLnexus/blob/v1.2.2/src/cli_utils.cc#L808-L852.

```
# Custom configuration for joint calling DeepVariant whole genome sequencing gVCFs.
unifier_config:
    min_AQ1: 10
    min_AQ2: 10
    min_GQ: 0
    monoallelic_sites_for_lost_alleles: true
genotyper_config:
    required_dp: 0
    revise_genotypes: true
    more_PL: true
    trim_uncalled_alleles: true
    liftover_fields:
        - orig_names: [MIN_DP, DP]
          name: DP
          description: '##FORMAT=<ID=DP,Number=1,Type=Integer,Description="Approximate read
depth (reads with MQ=255 or with bad mates are filtered)">'
          type: int
          combi_method: min
          number: basic
          count: 1
          ignore_non_variants: true
        - orig_names: [AD]
          name: AD
          description: '##FORMAT=<ID=AD,Number=R,Type=Integer,Description="Allelic depths
for the ref and alt alleles in the order listed">'
          type: int
          number: alleles
          combi_method: min
          default_type: zero
          count: 0
        - orig_names: [GQ]
          name: GQ
          description: '##FORMAT=<ID=GQ,Number=1,Type=Integer,Description="Genotype
Quality">'
          type: int
```

```
          number: basic
          combi_method: min
          count: 1
          ignore_non_variants: true
        - orig_names: [PL]
          name: PL
          description: '##FORMAT=<ID=PL,Number=G,Type=Integer,Description="Phred-scaled
genotype Likelihoods">'
          type: int
          number: genotype
          combi_method: missing
          count: 0
          ignore_non_variants: true
```

### Supplementary Note 2: GATK VQSR parameters

For running VQSR, we followed the settings used for the deep coverage 1KGP phase 3 release from NYGC (http://ftp.1000genomes.ebi.ac.uk/vol1/ftp/data_collections/1000G_2504_high_coverage/20190405_NYGC_b38_pipeline_description.pdf), but using GATK v4.1.2.0 instead of GATK v3.5 for our custom GIAB, CSER, and PAGE cohort. Note that some parameters in GATK v3.5 have different names in GATK4. All resource files can be found in Broad Institute's public resource directory on Google Cloud Storage (gs://genomics-public-data/resources/broad/hg38/v0).

```
RES_HAPMAP="hapmap_3.3.hg38.vcf.gz"
RES_1KG_OMNI="1000G_omni2.5.hg38.vcf.gz"
RES_1KG_P1_SNP="1000G_phase1.snps.high_confidence.hg38.vcf.gz"
RES_MILLS_INDEL="Mills_and_1000G_gold_standard.indels.hg38.vcf.gz"
RES_DBSNP="Homo_sapiens_assembly38.dbsnp138.vcf"

gatk --java-options -Xmx8g VariantRecalibrator
    -R "${ref_genome_fasta}"
    -V "${input_vcf}"
    -O "${output_snp_basepath}.recal"
    --tranches-file "${output_snp_basepath}.tranches"
    --rscript-file "${output_snp_basepath}.plots.R"
    -mode SNP
    "--resource:hapmap,known=false,training=true,truth=true,prior=15.0" "${RES_HAPMAP}"
    "--resource:omni,known=false,training=true,truth=true,prior=12.0" "${RES_1KG_OMNI}"
    "--resource:1000G,known=false,training=true,truth=false,prior=10.0" "${RES_1KG_P1_SNP}"
    "--resource:dbsnp,known=true,training=false,truth=false,prior=2.0" "${RES_DBSNP}"
    -an QD
    -an MQ
    -an FS
    -an MQRankSum
    -an ReadPosRankSum
    -an SOR
    -an DP
    -tranche 100.0
    -tranche 99.8
    -tranche 99.6
    -tranche 99.4
    -tranche 99.2
    -tranche 99.0
    -tranche 95.0
    -tranche 90.0
    --max-attempts 3
```

```
gatk --java-options -Xmx8g VariantRecalibrator
    -R "${ref_genome_fasta}"
    -V "${input_vcf}"
    -O "${output_indel_basepath}.recal"
    --tranches-file "${output_indel_basepath}.tranches"
    --rscript-file "${output_indel_basepath}.plots.R"
    -mode INDEL
    "--resource:mills,known=true,training=true,truth=true,prior=12.0" "${RES_MILLS_INDEL}"
    "--resource:dbsnp,known=true,training=false,truth=false,prior=2.0" "${RES_DBSNP}"
    -an QD
    -an FS
    -an ReadPosRankSum
    -an MQRankSum
    -an SOR
    -an DP
    -tranche 100.0
    -tranche 99.0
    -tranche 95.0
    -tranche 92.0
    -tranche 90.0
    --max-gaussians 4
    --max-attempts 3

gatk --java-options -Xmx8g ApplyVQSR
    -R "${ref_genome_fasta}"
    -V "${input_vcf}"
    -O "${vqsr_snp_vcf}"
    -mode SNP
    --truth-sensitivity-filter-level 99.80
    --recal-file "${output_snp_basepath}.recal"
    --tranches-file "${output_snp_basepath}.tranches"

gatk --java-options -Xmx8g ApplyVQSR
    -R "${ref_genome_fasta}"
    -V "${vqsr_snp_vcf}"
    -O "${vqsr_final_vcf}"
    -mode INDEL
    --truth-sensitivity-filter-level 99.0
    --recal-file "${output_indel_basepath}.recal"
    --tranches-file "${output_indel_basepath}.tranches"
```

### Supplementary Note 3: Reference panel creation

This is a Bash script for creating a reference panel from a 1KGP cohort VCF (DV-GLN-OPT or GATK-VQSR), closely following a standard pipeline in Eagle's website (https://data.broadinstitute.org/alkesgroup/Eagle/#x1-300005.3)

```
# Required tools: bcftools, tabix, Eagle.

# Input cohort VCF (from DeepVariant-GLnexus or GATK)
cohort_vcf="cohort-chr22.vcf.gz"

# 1KGP Reference genome:
ftp://ftp.1000genomes.ebi.ac.uk/vol1/ftp/technical/reference/GRCh38_reference_genome/GRCh38
_full_analysis_set_plus_decoy_hla.fa

ref_genome="GRCh38_full_analysis_set_plus_decoy_hla.fa"

# Genetic map file from Eagle repo:
https://data.broadinstitute.org/alkesgroup/Eagle/downloads/tables/genetic_map_hg38_withX.tx
t.gz

genetic_map_file="genetic_map_hg38_withX.txt.gz"

# Intermediate/output file names
cohort_processed_vcf="cohort-chr22-processed.bcf"
eagle_output_prefix="cohort-chr22-reference-panel"

# Filter singletons, apply variant filter, and convert to bcf.
bcftools view --no-version \
    -c 2 \
    -f ".,PASS" \
    "${cohort_vcf}" | \
  bcftools norm --no-version -Ou -m -any | \
  bcftools norm --no-version -Ob -o "${cohort_processed_vcf}" \
    -d none -f "${ref_genome}" && \
  bcftools index -f "${cohort_processed_vcf}"

# Run Eagle for phasing
eagle \
  --geneticMapFile=${genetic_map_file}" \
  --vcf="${cohort_processed_vcf}"
  --outPrefix="${eagle_output_prefix}" \
  --vcfOutFormat=z \
  --numThreads="$(nproc)"

# Index the output
tabix "${eagle_output_prefix}.vcf.gz"
```

### Supplementary Note 4: Genotype imputation

This is a Bash script for imputing phased (pseudo-)microarray variants with Beagle 5.0 using a reference panel.

```
# Required tools: Beagle 5.0, tabix

# Input files
reference_panel="cohort-chr22-reference-panel.vcf.gz"
input_vcf="HG002.pseudo-microarray.phased.chr22.vcf.gz"

# Output file name
output_prefix="HG002.imputed-pseudo-microarray.phased.chr22.vcf.gz"

chrom="chr22"

# Run Beagle for imputation
java -Xss2048k -Xmx50G -jar beagle.jar \
  "ref=${reference_panel}" \
  "gt=${input_vcf}" \
  "out=${output_prefix}" \
  "chrom=${chrom}"

tabix "${output_prefix}.vcf.gz"
```

### Supplementary Note 5: Software versions

DeepVariant (Google Brain): **v0.8.0** and **custom model** (included in 1000 Genomes data release) for NovaSeq reads.
GLnexus (DNAnexus): This study was run using **v1.2.0-pre.0**. The optimized parameters from this study are now included in GLnexus **v1.2.2** as two presets: `DeepVariantWGS` and `DeepVariantWES`.
GATK (Broad Institute): **v4.1.2.0**, except for the GATK 1KGP VCFs released by the New York Genome Center which used GATK **v3.5**.
Hap.py (Illumina): **v0.3.9**.
Eagle: **v2.4.1**.
Beagle: **v5.0**.
bcftools: **v1.9**.
